# Microcystin-RR is a biliary toxin selective for neonatal cholangiocytes

**DOI:** 10.1101/2023.08.09.552661

**Authors:** Kapish Gupta, Dongning Chen, Rebecca G. Wells

**Affiliations:** Division of Gastroenterology and Hepatology, Department of Medicine, University of Pennsylvania, Philadelphia, PA, USA; Center for Engineering MechanoBiology, University of Pennsylvania, Philadelphia, PA, USA; Department of Bioengineering, University of Pennsylvania, Philadelphia, PA, USA

**Author notes:** <u>Corresponding author:</u> Rebecca G. Wells 421 Curie Boulevard, 905 BRB II/III, Philadelphia, PA 19104. <u>Email:</u> Kapish Gupta Dongning Chen Rebecca G. Wells. <u>Author contributions</u> Kapish Gupta: Conceptualization, Formal analysis, Methodology, Investigation, Writing - original draft Dongning Chen: Methodology Rebecca G. Wells: Conceptualization, Resources, Funding acquisition, Supervision, Writing - review & editing.

**Keywords:** Algal toxins, biliary atresia, bile duct, glutathione and oxidative stress, harmful algal bloom

## Abstract

**BACKGROUND AND AIMS:** Biliary atresia is a fibrosing cholangiopathy affecting neonates that is thought to be caused by a prenatal environmental insult to the bile duct. Biliatresone, a plant toxin with an α-methylene ketone group, was previously implicated in toxin-induced biliary atresia in Australian livestock, but is found in a limited location and is highly unlikely to be a significant human toxin. We hypothesized that other molecules with α-methylene ketone groups, some with the potential for significant human exposure, might also be biliary toxins.

**APPROACH AND RESULTS:** We focused on the family of microcystins, cyclic peptide toxins from blue-green algae that have an α-methylene ketone group and are found worldwide, particularly during harmful algal blooms. We found that microcystin-RR, but not 6 other microcystins, caused damage to cell spheroids made using cholangiocytes isolated from 2-3-day-old mice, but not from adult mice. We also found that microcystin-RR caused occlusion of extrahepatic bile duct explants from 2-day-old mice, but not 18-day-old mice. Microcystin-RR caused elevated reactive oxygen species in neonatal cholangiocytes, and treatment with N-acetyl cysteine partially prevented microcystin-RR- induced lumen closure, suggesting a role for redox homeostasis in its mechanism of action.

**CONCLUSIONS:** This study highlights the potential for environmental toxins to cause neonatal biliary disease and identifies microcystin-RR acting via increased redox stress as a possible neonatal bile duct toxin.

## Introduction

Biliary atresia (BA) is a potentially fatal disease of neonates. It is a fibrosing cholangiopathy that is diagnosed within the first months of life but is thought to result from a prenatal environmental insult that damages the fetal extrahepatic bile duct (EHBD), causing duct obstruction and, ultimately, end-stage liver disease. BA occurs in 1:3,500 - 1:19,000 live births worldwide and is one of the most important pediatric liver diseases and the major indication for liver transplant in children in the United States [1–4]. Importantly, BA is never seen in older children or in the mothers of affected babies, suggesting a developmental susceptibility to the as-yet-unknown exposure [3]. The disease is characterized by progressive inflammation, fibrosis, and obstruction of the bile ducts, resulting in extrahepatic duct obliteration with impaired bile flow, cholestasis, and fibrosis [5]. Despite advances in medical and surgical management, care of patients with BA remains a significant challenge, and early diagnosis and intervention are critical to improving outcomes and preventing long-term complications [6,7].

Although candidate genetic modifiers have been identified, BA is not a genetic disease. Rather, epidemiological data suggest that prenatal environmental insults such as viral infections and toxin exposure are primary causes [8–11]. In spite of extensive study, however, only a small percentage of BA cases have been linked to viral infections, primarily Cytomegalovirus [12], while the insult in the majority of cases remains unknown.

We and colleagues previously identified a plant toxin, biliatresone, as a likely cause of a BA-like disease in Australian livestock [9]. Biliatresone is toxic to the extrahepatic bile ducts of larval zebrafish and to mammalian cholangiocytes *in-vitro* and can cause a BA-like phenotype in mouse pups [13–17]. Biliatresone is highly electrophilic and has an α-methylene ketone group that can form adducts with cysteine residues in proteins, affecting redox metabolism [18]. Several lines of evidence implicate redox stress in neonatal EHBD damage: the EHBDs of neonates have low levels of reduced glutathione (GSH) at baseline; artificially lowering GSH causes damage in cholangiocyte spheroids and neonatal EHBD explants; and N-acetyl cysteine treatment protects against damage from biliatresone [13,19].

While it is highly unlikely that pregnant women are exposed to biliatresone, these findings suggest that toxin exposure and redox stress during prenatal development could contribute to the development of BA. We hypothesized that environmental toxins that share a reactive group with biliatresone might also be biliary toxins. The 250+ toxins in the microcystin family, which are found worldwide, are particularly attractive candidates. Microcystins are cyclic heptapeptides with L-amino acid residues at positions 2 and 4 (Fig. 1A); microcystin-LR (leucine-arginine; MC-LR) is the most prominent congener and MC-RR (arginine-arginine) the second most commonly identified [20–22]. Like biliatresone, microcystins have an α-methylene ketone group. They are potent hepatotoxins produced by various species of cyanobacteria (often *Microcystis* species) that occur commonly as part of harmful algal blooms in freshwater ecosystems worldwide [23]. Microcystins are a significant threat to public health, accumulating in aquatic organisms such as fish and shellfish and entering the human food chain [24]. Additionally, exposure to microcystins can occur through recreational activities such as swimming and boating, as well as through drinking water contaminated with cyanobacteria [25].

**Figure 1:**
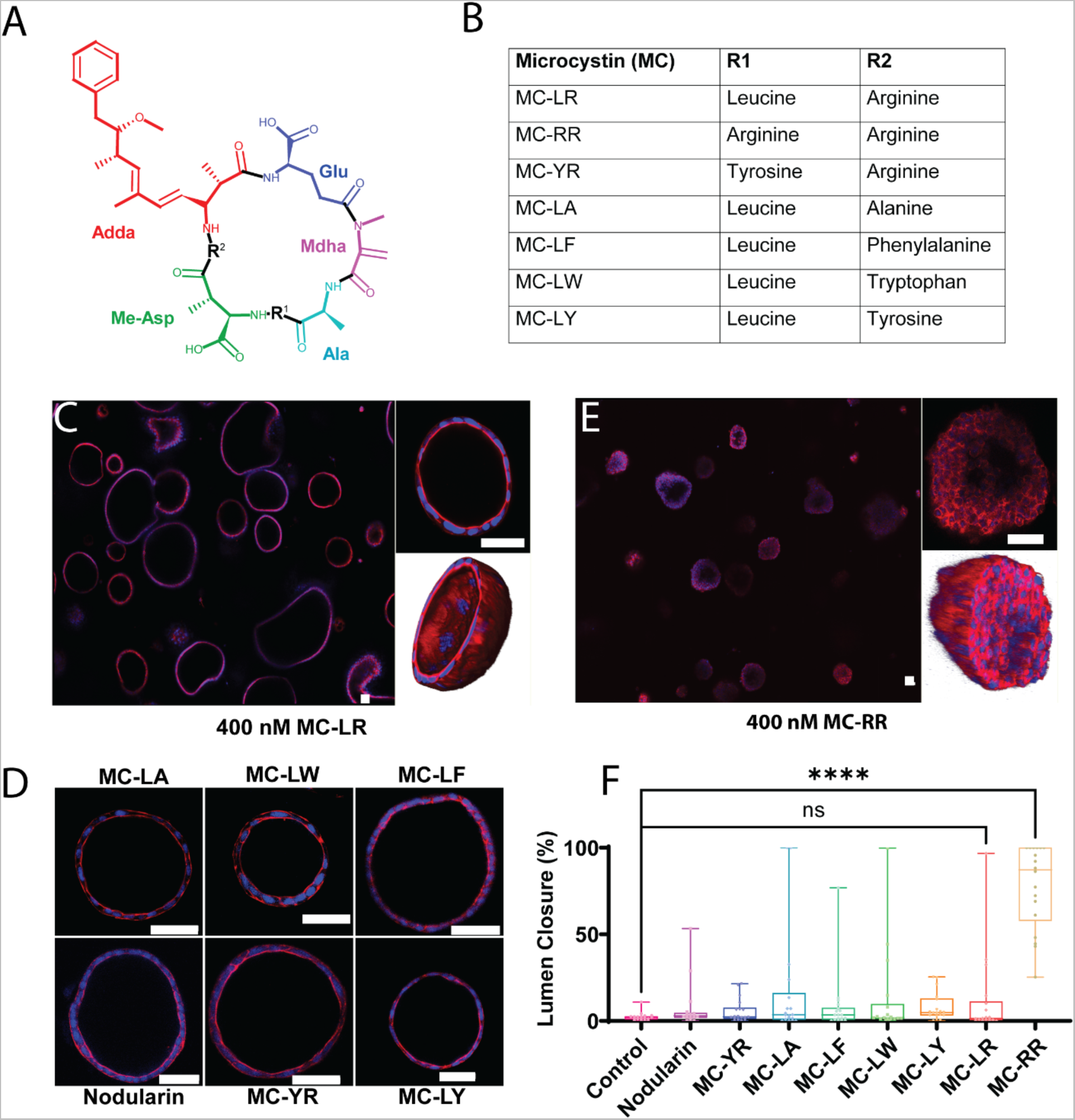
MC-RR treatment results in neonatal EHBD cholangiocyte spheroid damage. A) General structure of microcystins showing cyclic heptapeptide backbone with two variable amino acid groups (marked as R^1^ and R^2^). Mdha: N-methyldehydroalanine, Adda: (all-S,all-E)-3-amino-9-methoxy-2,6,8-trimethyl-10-phenyldeca-4,6-dienoic acid, Me-Asp: D-erythro-β-methyl-isoaspartic acid, Ala: Alanine, Glu: Glutamic acid. B). Commercially available MCs tested, with identity of amino acids at R^1^ and R^2^ noted. C) Neonatal EHBD cholangiocyte spheroids forming one-cell thick hollow spheroids in presence of 400 nM MC-LR. The middle section of a representative spheroid is highlighted and the 3D rendering shows a typical hollow lumen. D) Neonatal EHBD cholangiocyte spheroids form one-cell thick hollow spheroids with filled lumens after treatment with 400 nM of various microcystins and nodularin for 24 h. E) Neonatal EHBD cholangiocyte spheroids forming multi-cell thick spheroids after treatment with 400 nM MC-RR for 24 h. The middle section of a representative spheroid is highlighted and the 3-D rendering shows the lumen filled with cells. F) Quantification of lumen closure in the presence of all the tested microcystins and nodularin. All images are representative of 3 independent experiments, with a minimum of 18 spheroids for each condition quantified for lumen closure. Control spheroids are vehicle treated. Red: actin, Blue: DAPI. All scale bars 50 µm.

In this study, we evaluated the toxicity of seven commercially-available microcystins (Fig. 1B) and a related cyanobacterial toxin, nodularin, also with an α-methylene ketone moiety, on primary mouse and rat cholangiocytes and EHBD explants. We compared the effects of the different toxins on neonatal vs. adult cells and tissues, and identified a partial mechanism of action for MC-RR, the one microcystin found to be toxic in our systems. Our results suggest that MC-RR should be investigated as a cause of human BA.

## Methods

### Reagents

Microcystins were obtained from Enzo Life Sciences (Farmingdale, NY, USA) and Cayman Chemical Company (Ann Arbor, MI, USA). The Microcystin (Adda specific) ELISA kit was purchased from Enzo Life Sciences. Pranlukast (P0080), thiazolyl blue tetrazolium bromide (M2128), suramin sodium salt (S2671), and N-acetyl-L-cysteine (NAC) (A9165) were from Sigma (Burlington, MA, USA). Rat tail collagen (354236) and Matrigel (354234) were from Corning (Corning, NY, USA). The TUNEL assay kit (C10247) and CellRox Green (C10444) were obtained from Invitrogen (Waltham, MA, USA). A list of antibodies and dyes used in the study can be found in Supporting Table S1.

### Experimental animals

BALB/c mice (P2-3, P15-18, and adult) and Sprague Dawley rats (CD IGS, P2-3) were used in this study. Animals were euthanized and EHBDs were isolated and used either in cell culture or for direct experimentation as described in the Supporting Information. All animal experiments were performed in accordance with National Institutes of Health policy and were approved by the Institutional Animal Care and Use Committee of the University of Pennsylvania (Protocol number: 804862).

### Spheroid culture

Cholangiocytes were isolated by outgrowth from the EHBDs of neonatel (P2-3) or adult mice, as described in [26]. Isolated cells were either cultured on collagen-coated dishes or in 3D in a collagen-Matrigel mixture [15]. Cholangiocytes in the collagen-Matrigel mixture formed spheroids within 5 d and were used for experiments 6-8 d after plating. EHBDs in collagen and cholangiocytes were cultured in biliary epithelial cell (BEC) media as described in [26] with the exception that the final serum concentration was 10 percent. The media was changed every 2 d. During toxin treatment, the media was replaced with BEC media devoid of serum (referred as 0% BEC media). Following treatments, spheroids were fixed and stained as described in the Supporting Information.

### Explant Culture

Intact EHBDs from P2 mice, P15-18 mice or P2 rats were cultured at 37°C in 95% O_2_/5% CO_2_ in a Vitron Dynamic Organ Culture Incubator for 1 d in the presence of vehicle or a toxin. Ducts were stained as described previously [27] using antibodies against KRT19 and vimentin, and counterstained with DAPI (Supporting Information).

### Image Analysis

All samples were imaged using a water immersion 25X objective lens on a Leica TCS SP8 confocal microscope. Z-stacks (at a distance of 5 µm) were obtained for spheroids and EHBDs. Spheroids were analyzed using a custom-made algorithm as described in Supporting Fig. S1 using imageJ. EHBD images were stitched together and shown in this work with respect to base stack as described in Supporting Fig. S2. EHBDs were scored as normal, partially damaged, or completely damaged as per the scoring method described in Supporting Fig. S3.

### Statistical analysis

Statistical significance was calculated by either the one-tailed Student t-test or one-way analysis of variance followed by Tukey’s test for post-hoc analysis. P values ≤ 0.05 (*), p ≤ 0.01 (**), p ≤ 0.001 (***) and p ≤ 0.0001 (****) were considered statistically significant.

## Results

### Microcystin-RR damages neonatal cholangiocytes in spheroid culture

Cholangiocytes isolated from EHBDs can self-organize into spheroids with a hollow lumen (Supporting Fig. S4A). We observed that the morphology of spheroids generated from neonatal mouse cholangiocytes was unchanged compared to vehicle treated control spheroids in the presence of nodularin and most of the microcystins tested, indicating their low toxicity (Fig. 1C, D, F and Supporting Fig. S4E). However, exposure to 400 nM microcystin (MC)-RR resulted in loss of the spheroid monolayer and lumen closure (Fig. 1E & F), as assessed by a novel metric to quantitatively measure lumen closure; this was used for all lumen closure measurements throughout (Supporting Fig. S1). MC-RR- induced lumen closure was concentration-dependent, with higher concentrations resulting in greater damage (Fig. 2A & B). MC-RR exposure also caused a loss of polarity, as evidenced by a significant decrease in the ratio of apical to basal actin (Fig. 2C, Supporting Fig. S4). In contrast to these findings with neonatal EHBD cholangiocytes, spheroids derived from adult EHBD cholangiocytes showed no damage in the presence of MC-RR (Fig. 2D & E).

**Figure 2:**
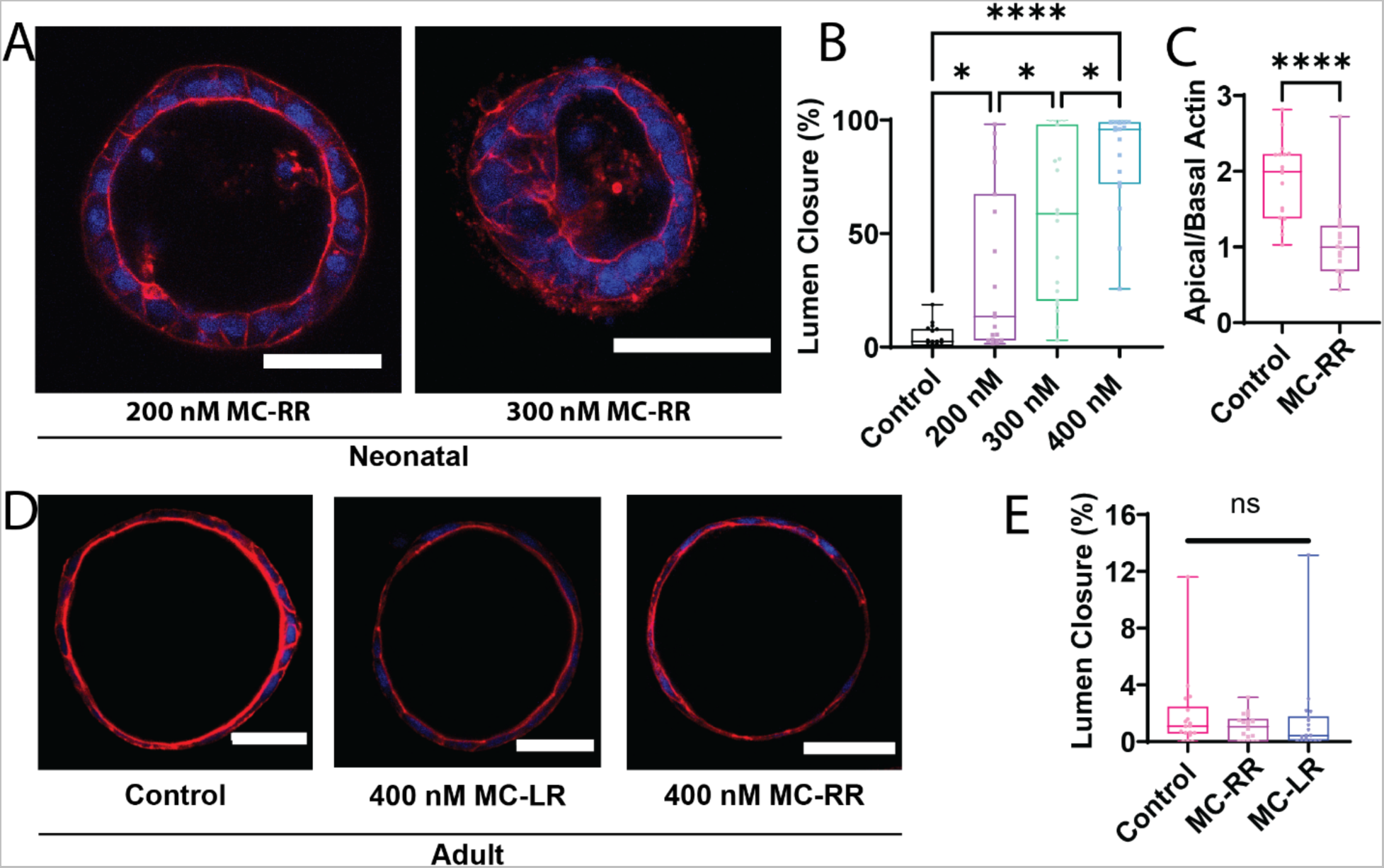
MC-RR-induced EHBD cholangiocyte spheroid collapse is specific for neonatal cholangiocytes. A) Degree of MC-RR-induced neonatal EHBD cholangiocyte spheroid collapse at varying MC-RR concentrations. B) Quantification showing dose dependence of lumen collapse. Representative images of spheroids treated with 200 and 300 nM MC-RR shown. Data for 400 nM MC-RR are from the experiment reported in Fig. 1. C) Quantification of loss of polarity (decrease in apical to basal actin ratio) in the presence of 400 nM MC-RR. D) Adult EHBD cholangiocyte spheroids show no damage after treatment with 400 nM of either MC-LR or MC-RR. E) Quantification showing no difference in lumen closure after treatment of adult EHBD cholangiocytes with either MC-LR or MC-RR. All images are representative of at least 3 independent experiments, with at least 18 spheroids quantified for lumen closure for each condition. Control spheroids are vehicle treated. Red: actin, Blue: DAPI. All scale bars 50 µm.

### MC-RR induces progressive and irreversible damage to neonatal EHBD cholangiocyte spheroids

The effect of 400 nM MC-RR on cholangiocyte spheroids was time-dependent, with progressively more damage observed over time, and significant damage detected after 15 h of exposure (Fig. 3A and B). The damage was irreversible: removal of MC-RR- containing media and subsequent incubation with normal media did not result in recovery of lumen damage. We tested washout at both early (15 h) and late (24 h) treatment times and found no recovery from the lumen damage (Fig. 3C and D, Supporting Fig. S5). However, we observed that spheroids treated for 15 h followed by washout (which had partial lumen closure) did not exhibit further lumen closure (Fig. 3C), indicating that persistent MC-RR exposure is necessary for progressive damage.

**Figure 3:**
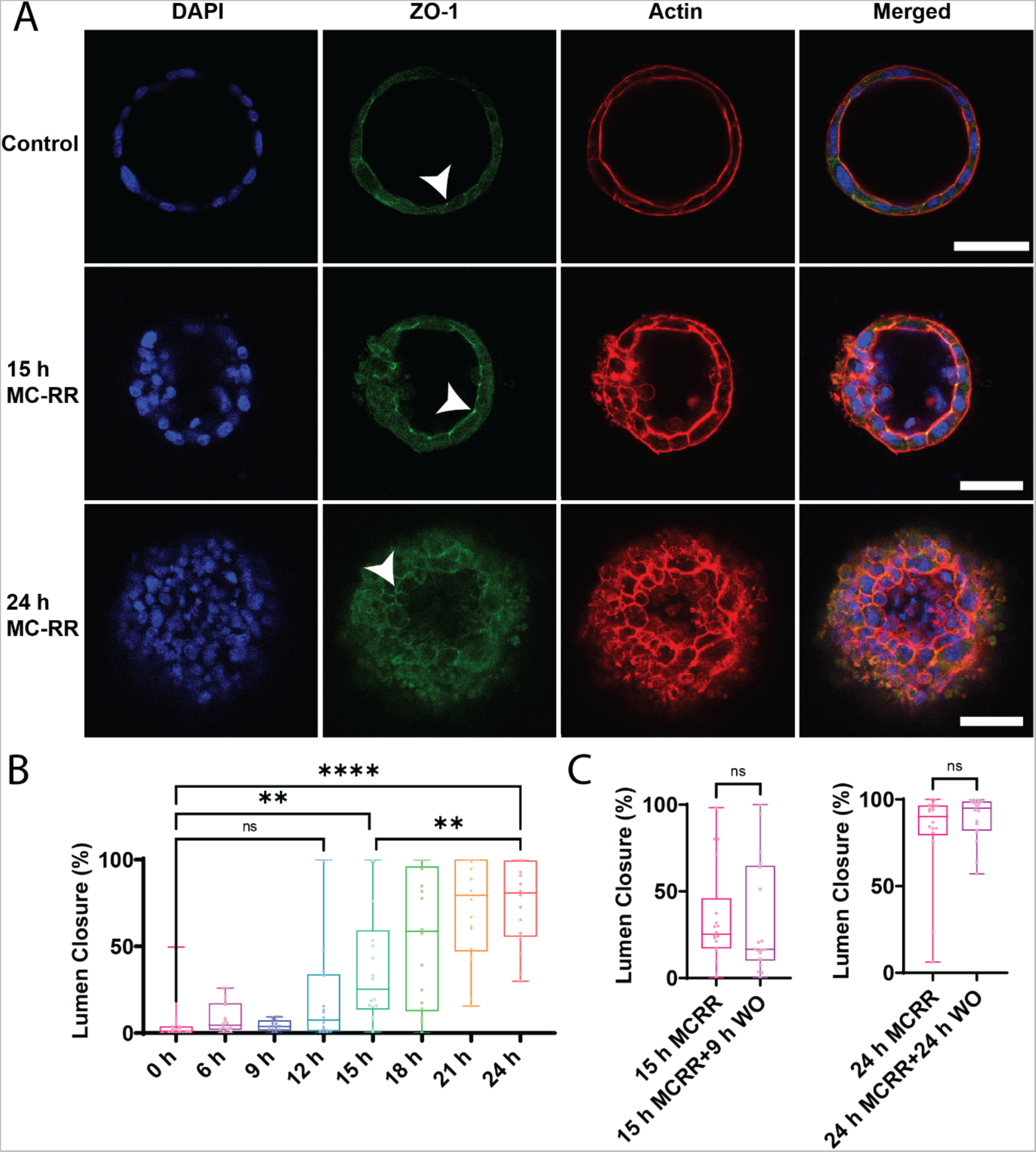
MC-RR induces progressive and irreversible lumen collapse. A) MC-RR (400 nm) induced increasing neonatal EHBD cholangiocyte spheroid collapse with increasing treatment time. Cell-cell contacts, as marked by ZO-1 staining, were maintained (white arrowhead) as the damage progressed. B) Quantification showing progressive lumen collapse with increasing MC-RR exposure time. C) Quantification showing that MC-RR-induced damage was irreversible after 15 h or 24 h exposure and that lumen patency did not improve even after MC-RR washout. All images are representative of at least 3 independent experiments, with at least 18 spheroids quantified for lumen closure. Control spheroids are vehicle treated. Red: actin, Green: ZO-1, Blue: DAPI. All scale bars 50 µm.

### MCRR results in damage of EHBD in explant culture

When mouse P2 EHBD explants were cultured in a high oxygen incubator in the presence of 400 mM MC-RR, we observed lumen closure, loss of monolayer integrity, and rounded cholangiocytes (Fig. 4A & B). Using a qualitative scoring method (described in Supporting Fig. S3), we found that these ducts exhibited significantly higher damage than ducts cultured with either the vehicle control or 400 nM MC-LR (the most common and most studied microcystin, used hereafter as a control) (Fig. 4C). o damage was observed in EHBD explants from P15-18 mice cultured with either MC-LR or MC-RR (Fig. 5A & B). Rat EHBDs showed similar results: EHBD explants from P2 pups showed severe damage after treatment with 400 nM MC-RR (Fig. 5C & D, Supporting Fig. S6). These results show that MC-RR induces significant and neonate-selective damage to EHBD explants. This damage was associated with increased vimentin staining in both neonatal EHBD explants and cholangiocyte spheroids (Supporting Figs. 7 A-C and S8).

**Figure 4:**
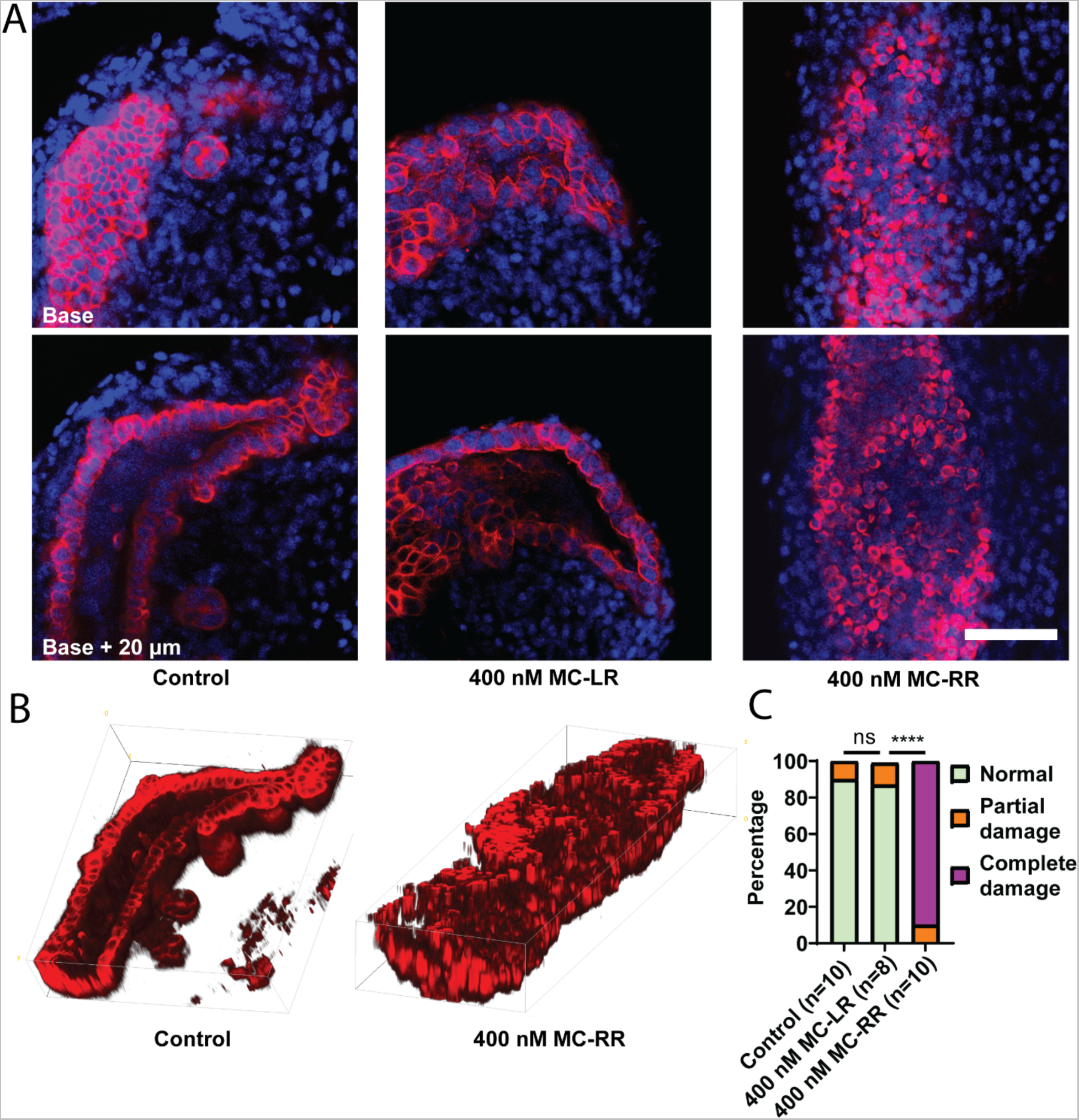
MC-RR treatment damages neonatal EHBD explants. A) Representative image showing neonatal EHBD explants cultured in a high oxygen incubator in the presence of vehicle, 400 nM MC-LR, or 400 nM MC-RR. EHBD explants were imaged using confocal microscopy; a base stack and a z-stack 20 µm above base are shown. B) 3D reconstruction of confocal images of EHBD explants treated with vehicle control and 400 nM MC-RR. C) Quantification showing significant EHBD damage in the presence of 400 nM MC-RR. The number of explants studied for each condition is shown in parentheses. Red: KRT19, Blue: DAPI. All scale bars 50 µm.

**Figure 5:**
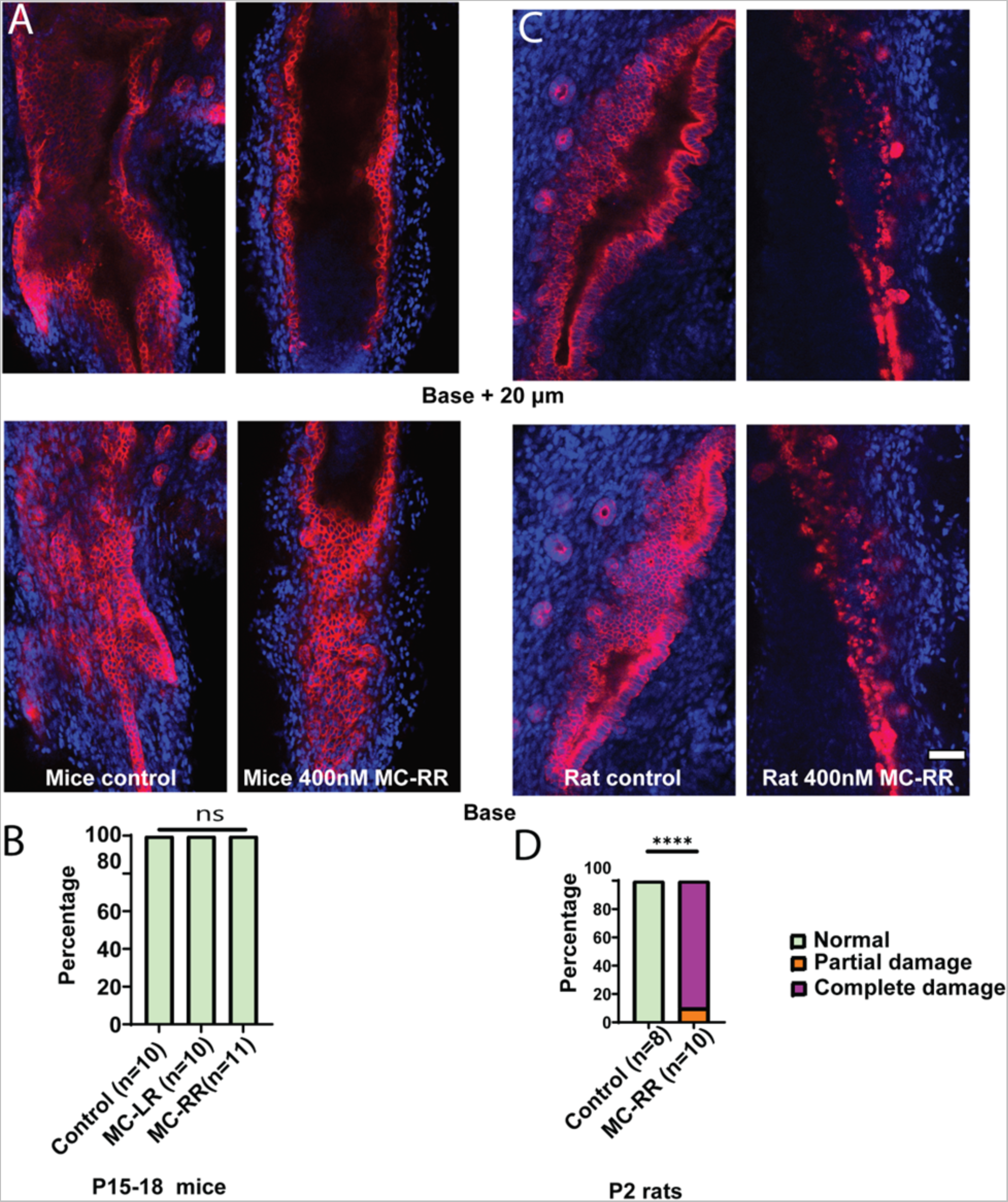
MC-RR specifically damages neonatal EHBDs. A) Representative images showing EHBDs isolated from P15-18 mice cultured as explants in the presence of vehicle or 400 nM MC-RR. B) Quantification showing lack of damage to EHBDs isolated from P15-18 mice pups. C) Representative image showing EHBDs isolated from P2 rat pups cultured as explants in the presence of vehicle or 400 nM MC-RR. D) Quantification showing damage to EHBDs isolated from P2 rat pups. The number of explants studied per condition is shown in parentheses. Red: KRT19, Blue: DAPI. All scale bars 50 µm.

### Mechanism of MC-RR-induced damage to neonatal cholangiocytes

Exposure to MC-RR did not cause apoptosis or cell proliferation of EHBD cholangiocytes, but instead resulted in the collapse of spheroids (Fig. 6A & B, Supporting Fig. S7D, E). In investigating the mechanism of MC-RR toxicity, we found that neonatal EHBD cholangiocytes took up significantly more MC-RR than MC-LR (Fig. 6C); however, uptake was similar for neonatal and adult cells (Fig. 6D). The observed differences in neonatal uptake were not due to differential activity of members of the organic anion transport protein (OATP) family. Neonatal EHBDs preferentially express OATP2a1 with little to no detectable expression of other OATP’s. Inhibiting OATP 2a1 using pranlukast or suramin [28] did not rescue cholangiocytes from MC-RR-induced damage (Supporting Fig. S9). Collectively, these data indicate that differential uptake of MC-RR is not the cause of neonatal cholangiocyte susceptibility to the toxin.

**Figure 6:**
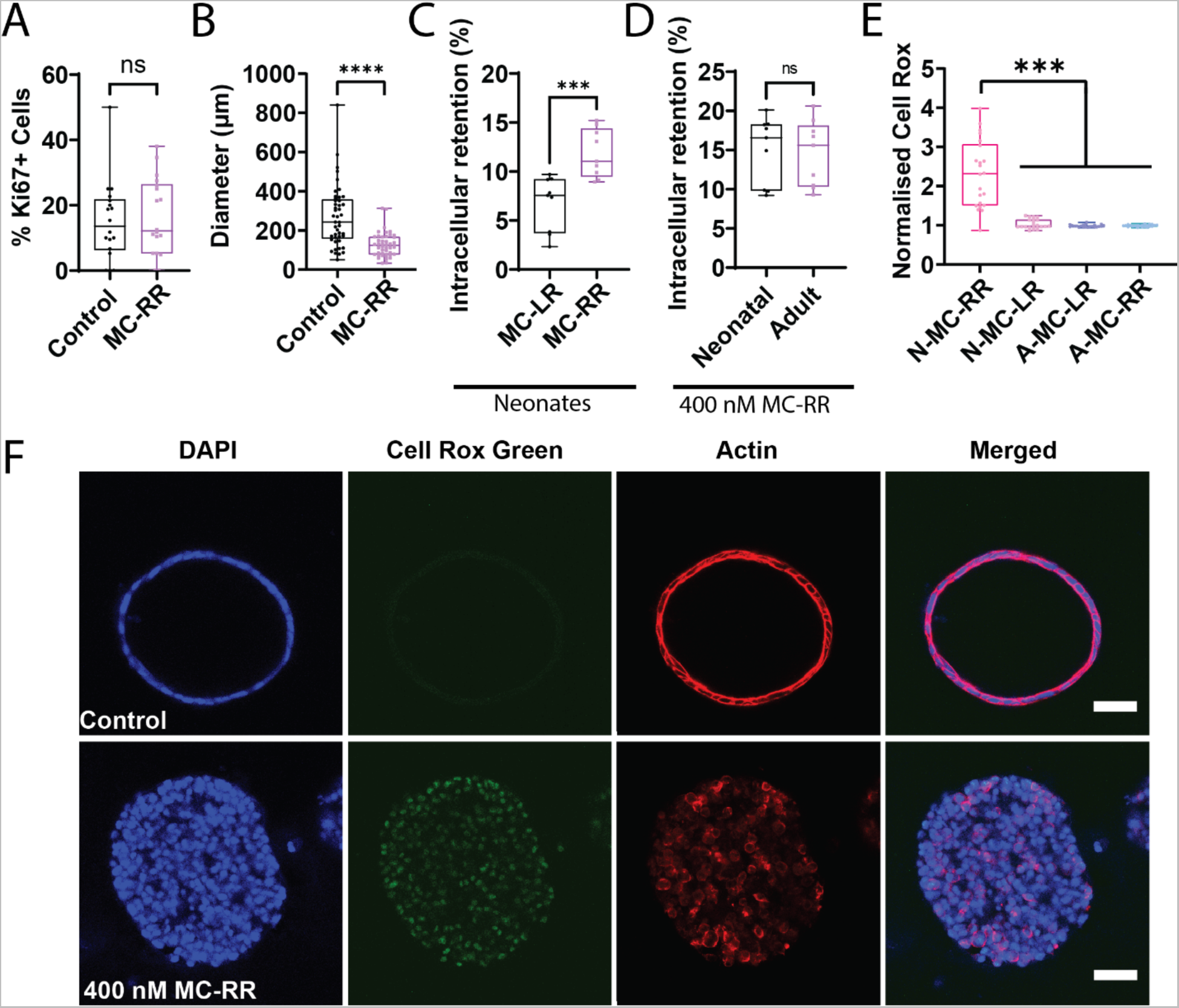
MC-RR induces elevated reactive oxygen species in neonatal EHBD cholangiocyte spheroids. A) Quantification of Ki67+ cells in neonatal mouse EHBD cholangiocyte spheroids treated with vehicle or 400 nM MC-RR (n=18 per condition from 3 independent experiments). B) Spheroid diameter after treatment with vehicle or 400 nM MC-RR (n=45 spheroids per condition from 3 independent experiments). C) Relative retention of MC-LR and MC-RR in neonatal EHBD cholangiocytes (n=9 samples from 3 independent experiments). D) MC-RR retention in neonatal and adult EHBD cholangiocytes (n=9 samples from each condition across 3 independent experiments). E) ROS production in neonatal and adult EHBD cholangiocyte spheroids after treatment with 400 nM MC-LR or 400 nM MC-RR for 24 h. F) ROS production in the presence of vehicle or 400 nM MC-RR. Images are representative of 3 independent experiments with at least 18 spheroids quantified for lumen closure per condition. Red: actin, Green: Cell-Rox Green, Blue: DAPI. All scale bars 50 µm.

We then considered the role of redox stress in MC-RR-induced cholangiocyte toxicity, and found that treatment with MC-RR induced higher levels of reactive oxygen species (ROS) in neonatal EHBD cholangiocyte spheroids compared to those treated with MC-LR. Additionally, ROS levels in MC-RR treated neonatal cholangiocytes were markedly higher than for adult EHBD cholangiocyte spheroids treated with either MC-LR or MC-RR (Fig. 6E & F, Supporting Fig. S10) and neonatal mouse EHBDs exhibit reduced levels of glutathione and various redox enzymes, including glutathione peroxidase (GPX)-1, GPX-4, catalase (CAT), and superoxide dismutase (SOD)-1 (Supporting Fig. S11). Collectively, these data suggest that differential susceptibility to redox stress in neonatal cholangiocytes [13,19,29] explains why they and not adult cholangiocytes are sensitive to MC-RR.

### N-Acetyl Cysteine rescues MC-RR-induced damage

We therefore hypothesized that treating neonatal EHBD cholangiocyte spheroids with N-acetylcysteine (NAC) would protect against damage from MC-RR. We found that treatment with NAC during exposure to 300 nM MC-RR resulted in partial blunting of the phenotype, as evidenced by maintenance of the hollow lumen and preservation of monolayer integrity (Fig. 7A). However, spheroids treated with NAC and 400 nM MC-RR exhibited a similar phenotype to those treated with 400 nM MC-RR alone, suggesting that the higher concentration of MC-RR may have overwhelmed the protective effects of NAC (Fig. 7B, Supporting Fig. S12). Similarly, in P2 mouse EHBD explant cultures, treatment with NAC coincident with 300 mM MC-RR led to a partial amelioration of the phenotype when compared to 300 nM MC-RR treatment, but no difference was observed with or without NAC when explants were treated with 400 nM MC-RR (Fig. 7C & D, Supporting Fig. S8).

**Figure 7:**
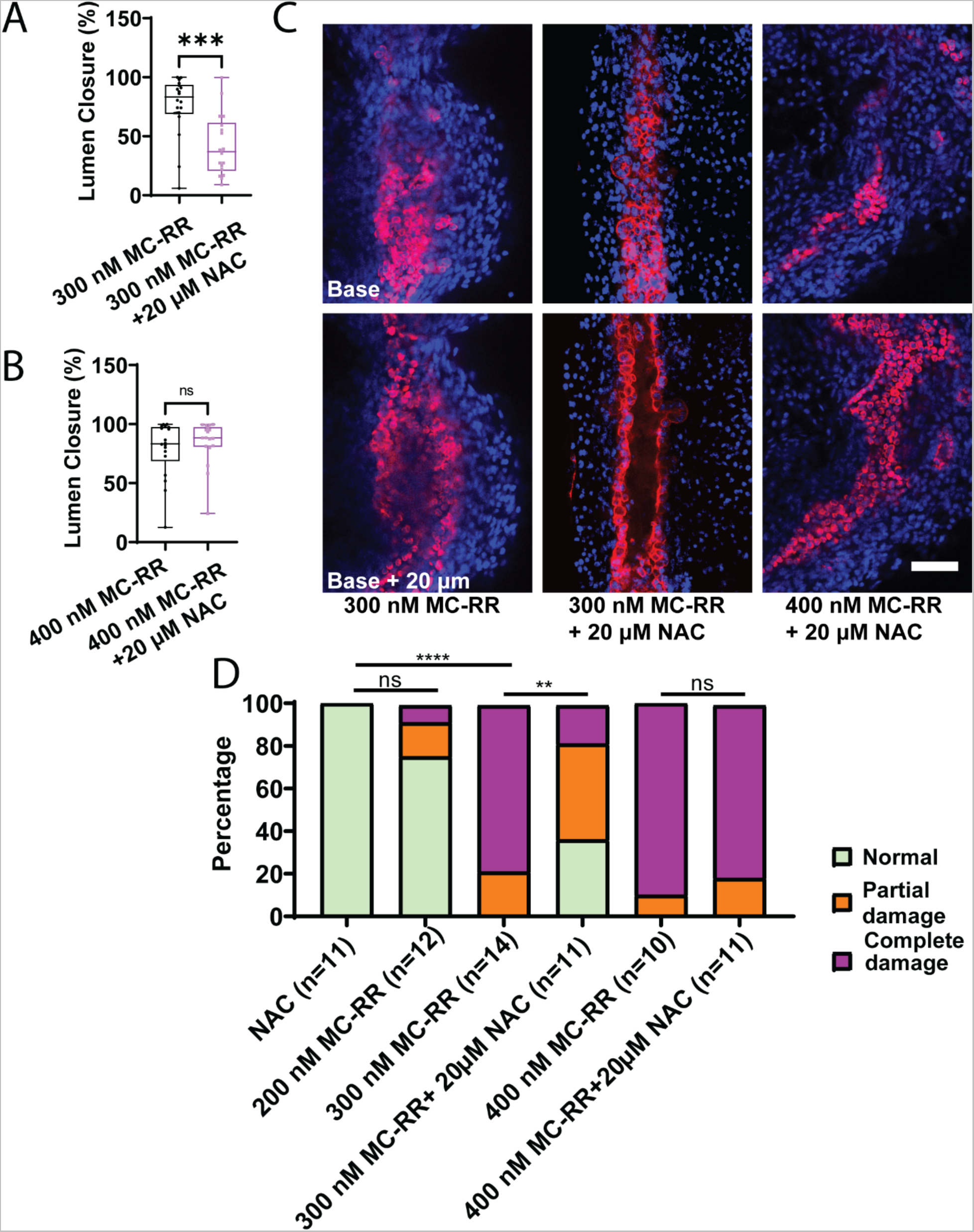
NAC partially ameliorates MC-RR induced damage in spheroids and EHBDs. A) Lumen closure in neonatal EHBD cholangiocyte spheroids treated with 300 nM MC-RR plus or minus NAC. B) Lumen closure in neonatal EHBD cholangiocyte spheroids treated with 400 nM MC-RR plus or minus NAC. C) Representative images showing MC-RR-induced EHBD damage at different MC-RR concentrations and in the presence of NAC. D) Quantification of EHBD explant damage after treatment with increasing MC-RR concentrations with and without NAC. Numbers of explants examined per condition are in parentheses. Red: KRT19, Blue: DAPI. All scale bars 50 µm.

## Discussion

Neonatal cholestatic diseases, including BA, are a leading cause of pediatric liver disease and pose significant treatment challenges [7]. While the cause of most cases of BA is unknown, recent research, including our own work, provides a proof-of-concept in animal models that toxin exposure of neonates or of the pregnant mother can cause a BA-like syndrome in neonates [14,16,17]. Toxins potentially responsible for BA in humans have not been identified – the toxin responsible for BA in neonatal livestock, biliatresone, is an unlikely source of toxin exposure for humans [9]. In this work, we examined the microcystins, a family of widely-distributed toxins with significance to humans and a structure including an α-methylene ketone, which is the active group in biliatresone [18,30]. We show here that one microcystin of seven tested, MC-RR, is toxic to rodent cholangiocytes and EHBD explants and notably that this toxicity is selective to neonatal, but not adult EHBDs. We further show that MC-RR specifically causes significant elevations in ROS in neonatal cholangiocytes and EHBD explants, a likely mechanism of action.

Microcystins are produced by cyanobacteria (blue-green algae), most in the genus *Microcystis*, during harmful algal blooms [24]. They are found globally and cause significant harm to humans and wildlife [23,31]; their impact is predicted to increase with climate change. The maximum safe value for MC-LR in drinking water, as established by the World Health Organization, is 1 µg/L (∼1nM) [32]. There are no specific guidelines for other microcystins and their toxicity is less well understood [32,33], but, of note, we were unable to locate reports of bile duct toxicity related to any of the microcystins.

MC-RR did not cause apoptosis or cell death, as determined by TUNEL staining and an MTT assay, but rather resulted irreversibly in a loss of apical-basal polarity and in architectural re-organization of the cholangiocyte monolayer in spheroids and EHBD explants. Increases in vimentin staining suggested a partial epithelial-to-mesenchymal transition [34], potentially aiding the cell migration or extrusion into the lumen that ultimately led to lumen obstruction. The epithelial-to-mesenchymal appeared to be at least partially independent of ROS, as vimentin expression also occurred in cholangiocytes following NAC treatment.

The specificity of MC-RR for neonatal, as opposed to adult, EHBD cholangiocytes and EHBDs was striking, particularly since MC-LR, which is more hepatotoxic than MC-RR [21], had no demonstrable effects. We found that treatment of cholangiocytes with MC-RR resulted in higher intracellular levels than treatment with MC-LR, although this was true for both neonatal and adult cells, and thus did not explain neonatal-specific toxicity. While microcystin uptake is often linked to OATPs, the literature does not provide a clear understanding of the specific OATP involved in MC-RR uptake in mice. Out of all the OATPs examined, the only OATP we found expressed on neonatal EHBD was OATP2a1, a prostaglandin transporter [35] with an unknown role in microcystin uptake. Treatment of spheroids with the known OATP2a1 inhibitors, pranlukast and suramin [28], had no effect on uptake, suggesting that MC-RR is taken up independently of OATP2a1. Microcystins target various organs such as intestine, liver, and kidney in mammals and, although they are mainly taken up in liver via OATPs, it is still not known how they are taken up by other organs such as kidney [23]. Many cell lines (including liver stem cell lines) that lack OATPs are also known to uptake microcystins [36], suggesting that there are other (currently unknown) transporters or cellular uptake mechanisms for microcystins.

The R^1^-position arginine of MC-RR appears to be necessary for toxicity, since neither MC-LR or MC-YR had any effect on cholangiocytes. Unfortunately, we were unable to test whether the R1 arginine was sufficient for toxicity. Although MC-RX compounds have been reported, none are available commercially and most have been identified only by LC-MS [30]; we were unable to locate any in sufficient amounts even through private sources. The presence of two arginine residues in MC-RR likely alters the microcystin ring structure. Understanding the role of the dibasic residues may be key to understanding MC-RR toxicity. Arginine residues in peptides can promote cellular uptake via direct translocation and endocytotic pathways [37]; however, whether the presence of two arginines in MC-RR promotes cellular uptake via this mechanism is not known.

The extrahepatic biliary system of neonates has lower levels of GSH than the intrahepatic system or adult ducts in either location, and has lower levels of key anti-oxidant pathway enzymes. This appears to play a key role in biliatresone toxicity [13,19,29]. Our finding that MC-RR induces higher levels of ROS in the neonatal EHBD is consistent with lower levels of GSH in the EHBD and with a role for oxidative stress in the mechanism of action of MC-RR. Recently reported research based on RNA sequencing of samples obtained during hepatoportoenterostomy surgery for BA suggests that a ROS-related mechanism may play a role in BA: a significant fraction of BA patients, oxidative pathways were found to be disturbed [38]. The effects of NAC, which replenishes cellular GSH and also directly interacts with and neutralizes ROS, including superoxide anions and hydroxyl radicals, raise the possibility that it could be used as a preventive or therapeutic agent [39] and indeed there is an ongoing clinical trial testing whether NAC has efficacy in BA [40]. Given that NAC only partially prevented damage in our rodent model systems, however, mechanisms in addition to redox stress may be at play.

We investigated microcystins due to their structural similarity to biliatresone. The damage induced by MC-RR exhibited similarities to that due to biliatresone in terms of spheroid damage and the preventative properties of NAC. However, there were notable differences between the effects of the two compounds as well. Biliatresone-induced damage required continuous exposure to the compound, with spheroids recovering after the removal of biliatresone (at least within a certain time frame [15]) while MC-RR-treated spheroids did not show recovery after drug washout. It is also important to note that the presence of an α-methylene ketone group in and of itself does not necessarily render a compound toxic. This is evident from the fact that other microcystins, which also contain this group, have no visible effect on cholangiocytes. These findings suggest that the mechanism underlying toxin-induced cholangiocyte damage is complex and warrants further investigation.

The occurrence of harmful algal blooms is on the rise and is projected to further increase as the climate continues to deteriorate [41]. Human exposure to microcystins will likely increase in parallel. In the United States, the incidence of BA cases has shown an upward trend, increasing from 28.5 cases per million births in 1997 to 55.5 cases per million births in 2012 [42]. Although BA cases do not exhibit clear seasonal patterns [43,44], there have been certain periods in which an increase in BA cases has been observed [45–49]. However, the current evidence is insufficient to establish a direct connection between these BA case increases and exposure to MC-RR. To ascertain any potential correlation between maternal exposure to MC-RR and the incidence of BA, further mechanistic studies and epidemiological investigations are needed. While MC-LR has been extensively studied, exploring the relationship between MC-RR and BA will require more detailed studies of MC-RR, as opposed to MC-LR, in bodies of water and in the food chain.

## Acknowledgements

We are grateful to the UPenn Cell and Developmental Biology Microscopy Core and the NIDDK Center for Molecular Studies in Digestive and Liver Diseases (P30 DK050306).

## List of Abbreviations

BA: Biliary atresia
BEC: Biliary epithelial cell
CAT: Catalase
EHBD: Extrahepatic bile duct
GSH: Glutathione
GPX-1: Glutathione peroxidase
IACUC: Institutional Animal Care and Use Committee
MC-LR: Microcystin-LR
MC-RR: Microcystin-RR
NAC: N-acetyl cysteine
OATP: Organic anion transport protein
ROS: Reactive oxygen species
SOD-1: Superoxide dismutase 1

## Supplemental Materials

### Supporting Material and Methods

#### Treatment of cholangiocyte monolayers

Cholangiocytes were cultured on collagen-coated dishes and treated with various concentrations of microcystins dissolved in 0% BEC media. Cell viability was assessed using the MTT assay, and apoptosis was measured using TUNEL staining. Additionally, microcystin retention was evaluated by treating cells with 400 nM MC-RR and MC-LR for 24 h, followed by washing with 1X PBS three times and incubation with 1X RIPA buffer for 30 min. The amount of microcystin in the cell lysate was then quantified using a Microcystin (Adda specific) ELISA kit, following the manufacturer’s instructions (Prod. No. ALX-850-319, Enzo Life Science Inc. Farmingdale, NY, United States).

#### Treatment of spheroids

Cholangiocytes were plated in a collagen-Matrigel mixture [1] and allowed to form spheroids for 6-8 days. These spheroids were then treated with 400 nM individual microcystins, 400 nM nodularin, or vehicle in 0% BEC media. To perform washout experiments, spheroids were treated with 400 nM MC-RR or vehicle control in 0% BEC media for 15 or 24 h, washed three times with 1X PBS, and then placed in 0% BEC media for 9 or 24 h, respectively. For rescue experiments, spheroids were treated with 400 nM MC-LR or MC-RR and drugs (20 µM NAC, 500 nM suramin, or 2 µM pranlukast [2] obtained from Invitrogen, Waltham, MA, USA) in 0% BEC media for 24 h. Subsequently, spheroids were fixed, stained, and imaged using a confocal microscope to evaluate spheroid morphology.

#### EHBD explant treatment

EHBD explants were treated with individual microcystins or vehicle in 0% BEC media in a Vitron Dynamic Organ Culture Incubator (San Jose, United States) for 24 h as described in [3]. For rescue experiments, EHBD were treated for 24 h with 400 nM MC-RR or 400nM MC-RR and 20 µM NAC in 0% BEC media. Following treatment, all EHBDs were stained as described below.

#### Immunofluorescence staining

Spheroids and EHBDs were fixed with paraformaldehyde, permeabilized using 0.5% Triton X-100 in PBS, and subsequently blocked with a blocking solution containing 10% FBS and 0.1% Triton X-100 in PBS. Immunostaining was performed as described previously (3) using antibodies listed in Table S1.

#### Quantitative real-time PCR

EHBDs from P2 (6 ducts per sample) and adult (3 ducts per sample) mice were homogenized using a Bullet Blender® Homogenizer and RNA was isolated using TRIzol™ Reagent (Invitrogen) according to the manufacturer’s instructions. 3 pooled samples each for P2 and adults were used. RNA concentrations were measured with a NanoDrop ND-1000 full-spectrum spectrophotometer (ThermoFisher Scientific, Wilmington, MA, USA). Single-stranded complementary DNA (cDNA) was synthesized using the SuperScript VILO cDNA Synthesis Kit (ThermoFisher) according to the manufacturer’s instructions. cDNA was diluted to 2 ng/μL for real-time quantitative PCR analysis and PCR reactions were performed using Taqman Mastermix (Applied Biosystems, ThermoFisher Scientific, Waltham, MA, USA) in a total volume of 10 μL. Real-time quantitative PCR was done for genes involved in oxidative response and for Organic anion transport protein (OATP) genes (also known as solute carrier organic anion transporters or Slco). The following predesigned primers were purchased from ThermoFisher: Sod (Mm01344233_g1), Gpx-1 (Mm00656767_g1), Cat (Mm00437992_m1), Gpx-4 (Mm04411498_m1), Gsto-1(Mm00599866_m1), Slco1a1 (Mm01267415_m1), Slco1a6 (Mm01267368_m1), Slco2a1 (Mm00459638_m1), Slco1a4 (Mm01267407_m1), Slco1b2 (Mm00451510_m1) and Slco2b1 (Mm00614448_m1). Thermal cycling and fluorescence detection were performed on a StepOnePlus Real-Time PCR system from ThermoFisher. Expression levels were corrected using a reference gene (ΔCt). For OATP genes, only 2a1 and 2b1 showed detectable expression (2^−ΔCt^) in both P2 and adult samples. All the genes involved in oxidative responses showed detectable expression. All PCR results results are shown as expression relative to expression in adults (2^−ΔCt_Sample^/2^−ΔCt_Adults^).

#### Glutathione (GSH) measurements

EHBD from P2 (4 ducts per sample) and adult (2 duct per sample) mice were homogenized using a Bullet Blender® Homogenizer in the presence of RIPA buffer (Invitrogen) and protease inhibitors. 6 pooled samples each for P2 and adults were used. Samples were centrifuges to remove debris. GSH was determined using Mouse GSH ELISA Kit (MyBioSource, Inc. San Diego, CA, USA) and protein content was determined using a micro BCA protein assay reagent kit (ThermoFisher Scientific). GSH per mg protein was determined for each sample. The results are shown as expression relative to expression in adults (GSH_Sample_/GSH_|Adult|_).

#### ROS measurements

Spheroids were treated with DMSO, 400 nM MC-LR, or 400 nM MC-RR for 24 h. After washing in 1X PBS, the spheroids were incubated with 5 µM CellRox Green for 30 min at 37°C. The samples were washed and counterstained with phalloidin and DAPI according to the manufacturer’s instructions, and then imaged.

---

**Supporting Table S1:** List of antibodies and stains.

| **Primary Antibodies** |  |  |  |  |
| --- | --- | --- | --- | --- |
| **Antigen** | **Host** | **Supplier** | **Cat. No.** | **Dilution** |
| KRT19 | Rabbit | Abcam, Boston, MA, USA | ab52625 | 1:100 |
| Vimentin | Chicken | Novus Biologicals, Centennial, CO USA | NB300-223 | 1:100 |
| Ki67 | Rabbit | Abcam | Ab16667 | 1:100 |
| ZO-1 | Rabbit | Invitrogen | 61-7300 | 1:50 |
| **Secondary Antibodies** |  |  |  |  |
| **Antigen** | **Host** | **Supplier** | **Cat. No.** | **Dilution** |
| Rabbit | Donkey | Jackson ImmunoResearch Laboratories, Inc, West Grove, PA, USA | 711-165-152 | 1:300 |
| Chicken | Donkey | Jackson ImmunoResearch Laboratories, Inc | 703-605-155 | 1:300 |
| **Stains** |  |  |  |  |
| **Dye** | **Target** | **Supplier** | **Cat. No.** | **Concentration** |
| DAPI | Nucleus | Invitrogen | D1306 | 1 µg/mL |
| Rhodamine Phalloidin | Actin | Invitrogen | R415 | 0.33 µM |
| CellRox Green | ROS | Invitrogen | C10444 | 5 µM |

## List of Supporting Figures

**Supporting Figure S1:**
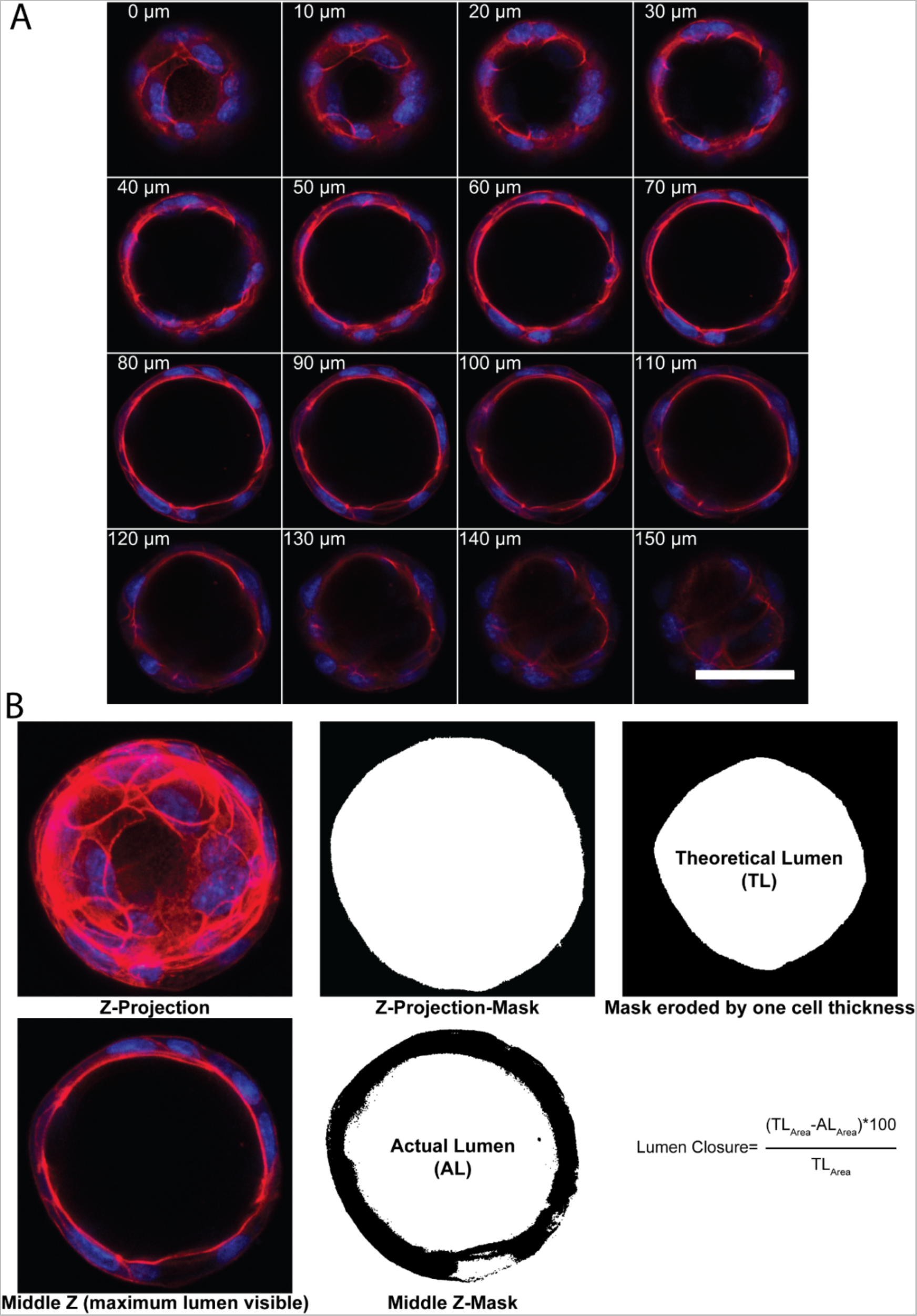
Quantification of lumen closure in EHBD cholangiocyte spheroids. A) Z-stacks (step size 5 µm) were obtained using confocal microscopy. Image shows all the slices for a representative spheroid. B) To quantify lumen closure, a Z-projection image was created and used to create a spheroid mask. The mask was eroded by 1 cell thickness (calculated by taking the average of the shortest diameter or width of 3 cells) to obtain theoretical lumen (TL) area, assuming a cell layer one cell thick. To calculate the actual lumen (AL) area, the middle Z-stack was used to create a mask. The relative difference in theoretical and actual lumen area was used to calculate “% lumen closure”. Red: actin, Blue: DAPI. All scale bars 50 µm.

**Supporting Figure S2:**
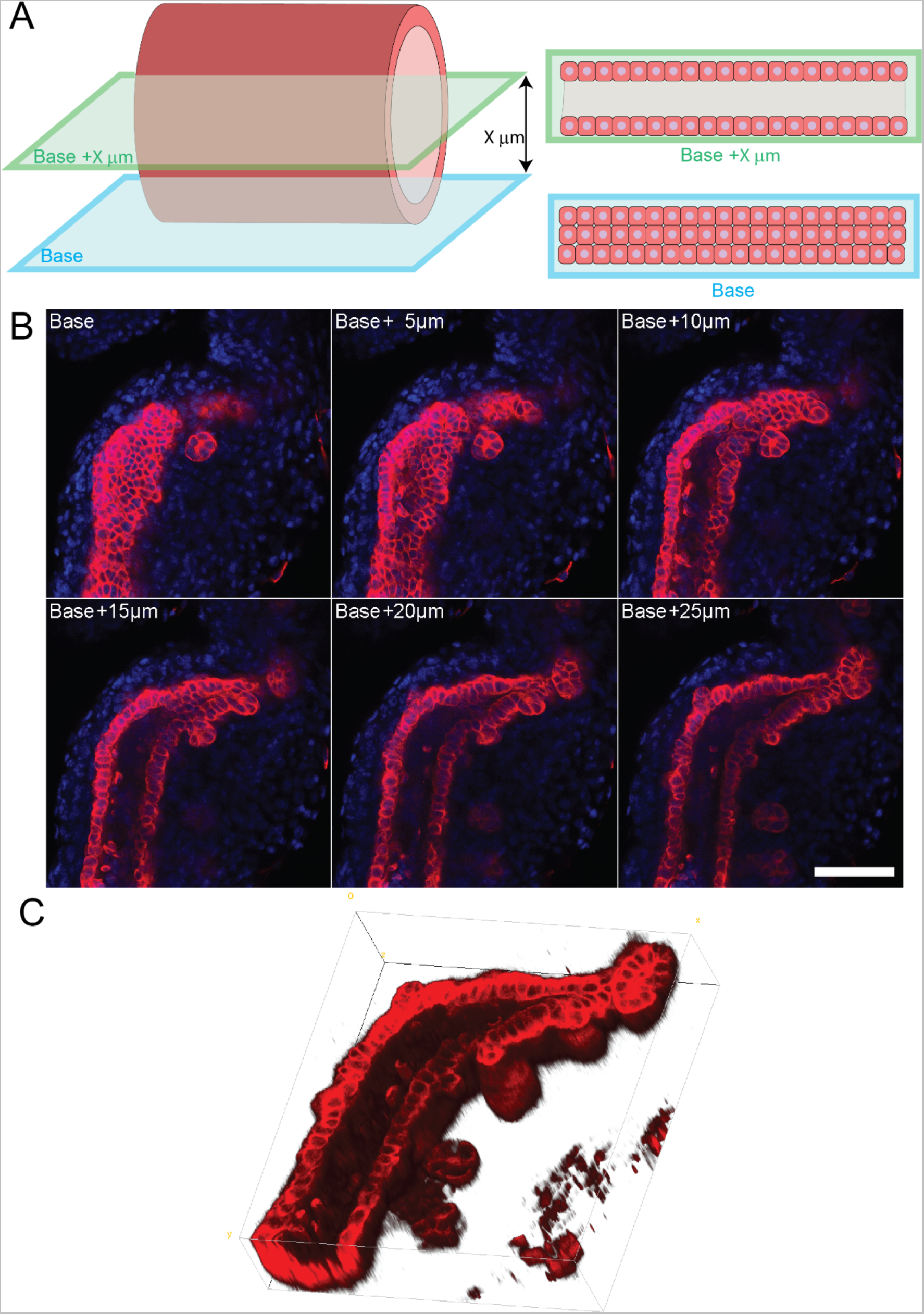
Imaging and representation of EHBD: A) Explant culture of EHBD in high oxygen incubator, without treatment. EHBDs were stained and imaged with confocal microscopy. Images were captured at 5 µm intervals relative to the “base” image, where the monolayer was visible. The images are displayed with respect to this base image, and each z-slice was annotated based on its distance from the base. The EHBD schematic depicts two sections: one at the base and another at X µm distance from the base. B) Representative image showing all the z-slices obtained for one EHBD. In Figure 4 only the “base” and “base+20 µm” slices are shown. C) 3-D rendering of the EHBD shown in B. Red: KRT19, Blue: DAPI. Scale bar 50 µm.

**Supporting Figure S3:**
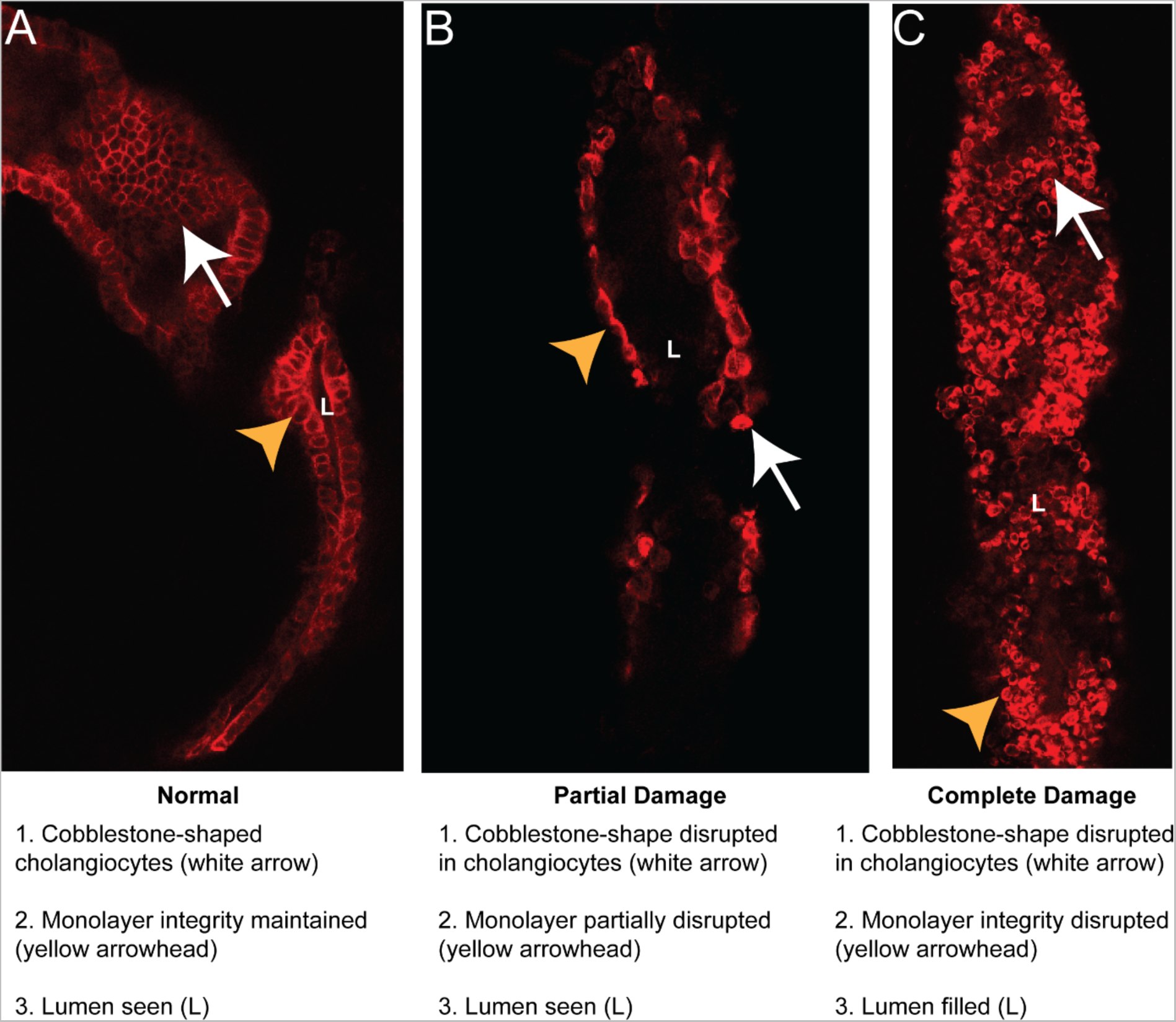
EHBD damage quantification. EHBDs were placed in one of three groups depending on the degree of damage, as assessed after KRT19 staining (red). A) EHBDs that showed cholangiocytes with a cobblestone morphology (white arrow) and forming a continuous monolayer (orange arrowhead) surrounding the lumen (L) were considered “normal”. B) EHBDs with abnormally-shaped cholangiocytes (white arrow) and disrupted monolayers (orange arrowhead) but with open lumens (L) were placed in the “partial damage” group. C) EHBDs with misshapen cholangiocytes (white arrow), a disrupted monolayer (orange arrowhead), and a closed lumen (L) were placed in the “complete damage” group. Red: KRT19. Scale bar 50 µm.

**Supporting Figure S4:**
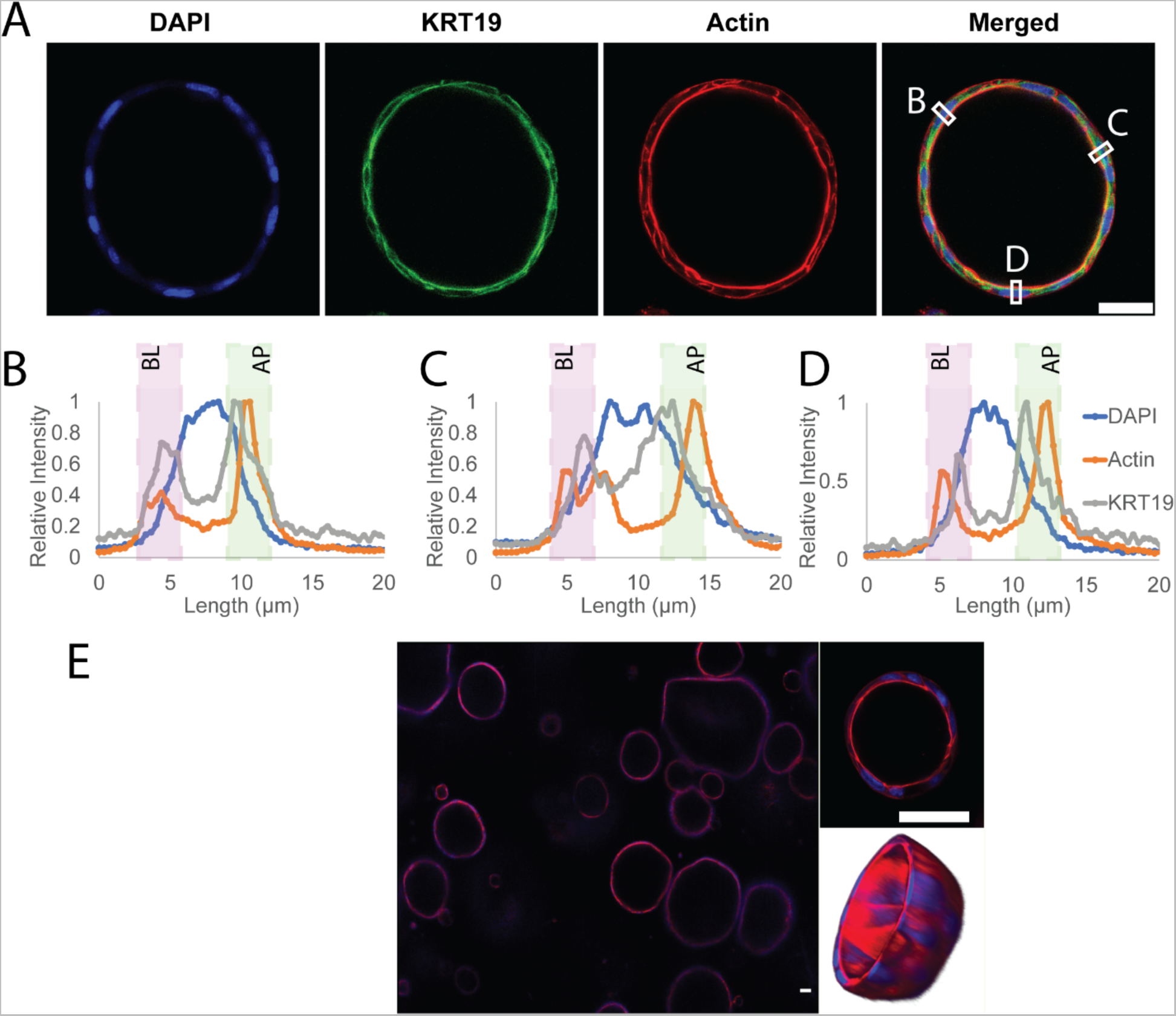
Neonatal EHBD cholangiocytes form hollow spheroids with apical/basal polarity. A) A control (untreated) spheroid stained with actin and KRT19. B-D) Intensity scans across lines marked as B, C and D in merged panel in A. Basal (BL) and apical (AP) regions are marked with purple and green rectangles. The cells have preferential apical actin localization and express more KRT19 apically. Nuclei are central. Ratio of maximum actin intensity in apical and basal sections, as determined by this method, was used in the quantification for main Fig. 2C. E) Representative image showing a cross-section of cholangiocytes cultured in a collagen-Matrigel mixture for 6 d and treated with vehicle for 24 h. Cells arrange themselves into hollow spheroids as shown in the 3D reconstruction. Red: actin, Green: KRT19, Blue: DAPI; all scale bars 50 µm.

**Supporting Figure S5:**
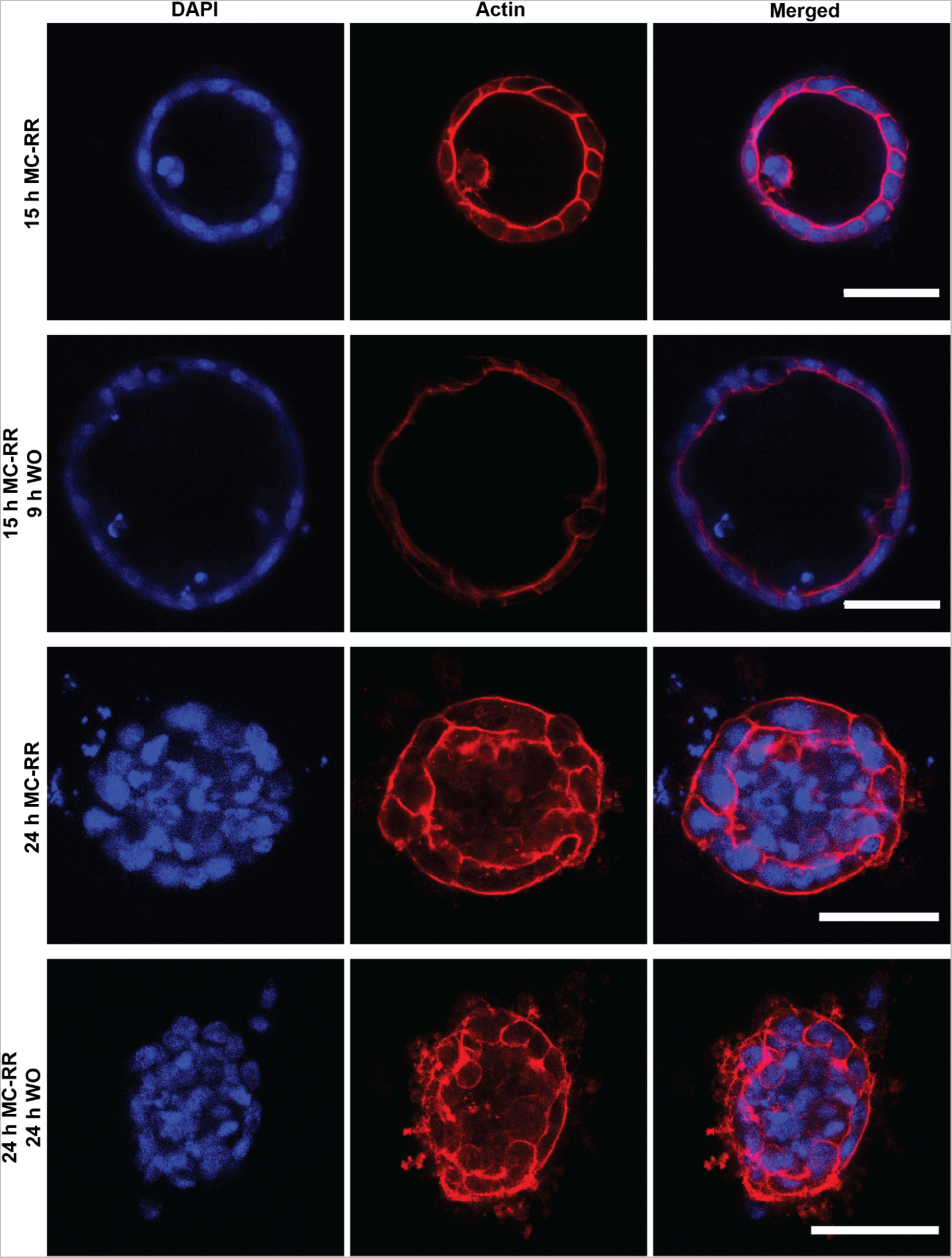
Washout after MC-RR treatment does not rescue MC-RR-induced damage. Representative images (similar to those in main Fig. 3) showing spheroids treated with 400 nM MC-RR for 15 and 24 h, with washout for 9 and 24 h, respectively. In both cases, removal of MC-RR with washout was not sufficient for rescue. Quantification in main Fig. 3. Red: actin, Blue: DAPI. All scale bars 50 µm.

**Supporting Figure S6:**
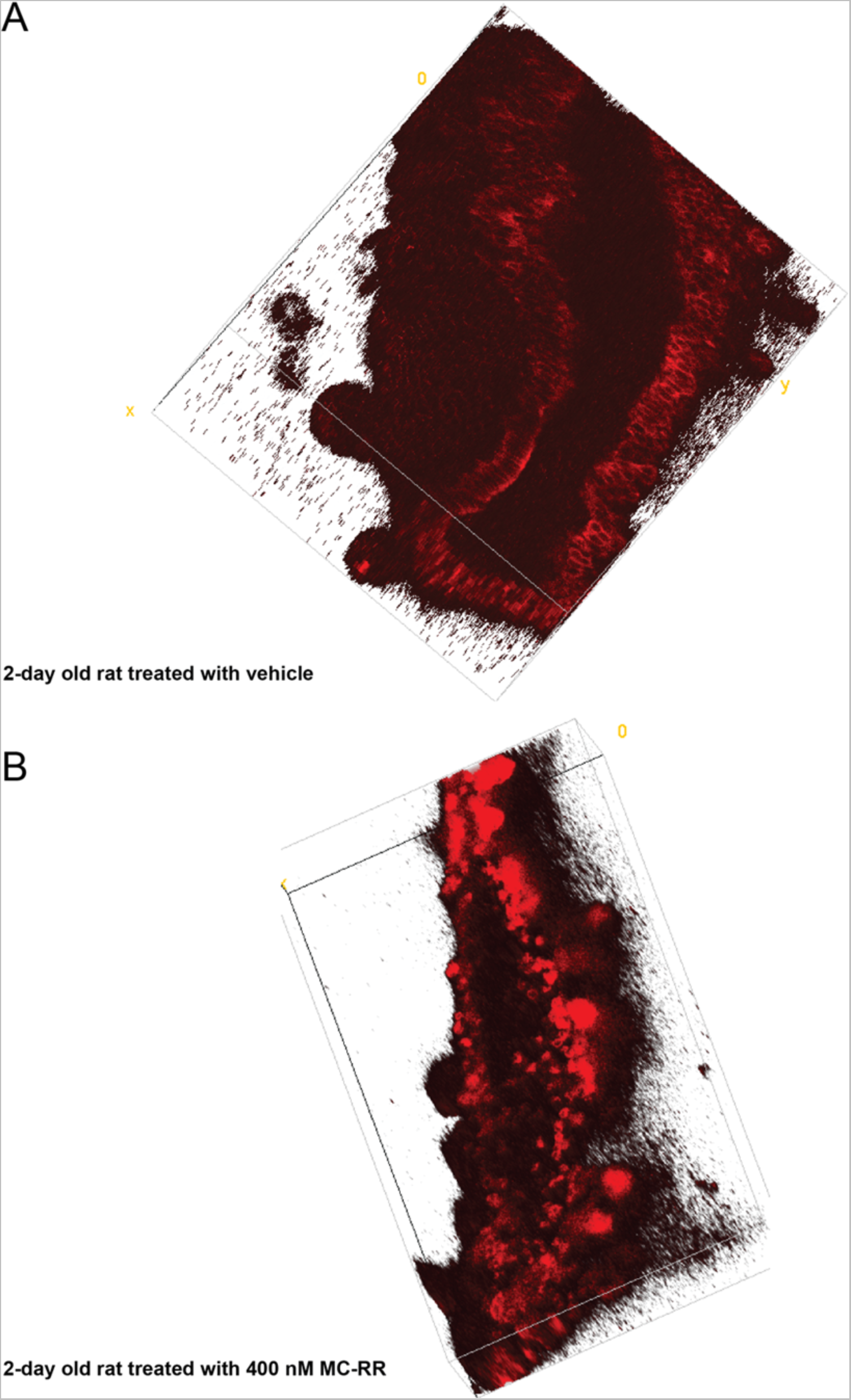
MC-RR-induced damage to neonatal rat EHBD explants. A) 3D reconstruction of representative EHBD isolated from P2 rat and cultured as explant in a high oxygen incubator. B) 3D reconstruction of representative EHBD isolated from P2 rat cultured similarly but in the presence of 400 nM MC-RR for 24 h. Red: KRT19.

**Supporting Figure S7:**
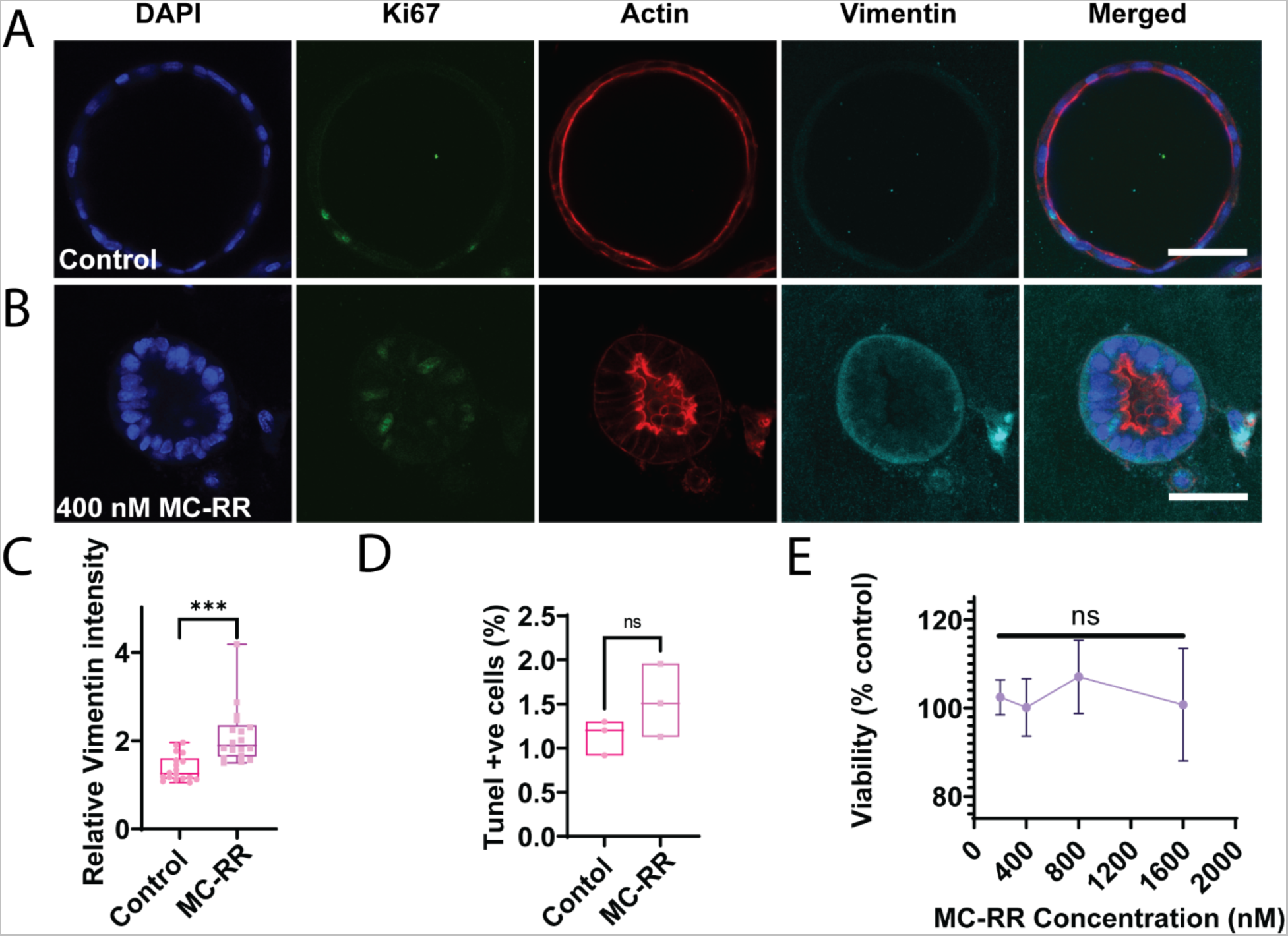
Mechanism of MC-RR-induced damage in neonatal mouse EHBD cholangiocyte spheroids. A) Representative image of control spheroids (treated with vehicle) and stained for Ki67 and vimentin. B) Representative image showing MC-RR-treated spheroids stained for Ki67 and vimentin. C) Quantification of vimentin staining intensity in EHBD cholangiocyte spheroids after treatment with 400 nM MC-RR for 24 h with a minimum of 18 spheroids for each condition quantified. D) TUNEL positive cells were measured after treatment with vehicle or 400 nM MC-RR for 24 h. Quantification from three independent experiments. E) MTT assay after treatment of cholangiocytes with MC-RR at concentrations up to 1.6 mM. Quantification from three independent experiments. Cyan: vimentin, Red: actin, Green: Ki67, Blue: DAPI. All scale bars 50 µm.

**Supporting Figure S8:**
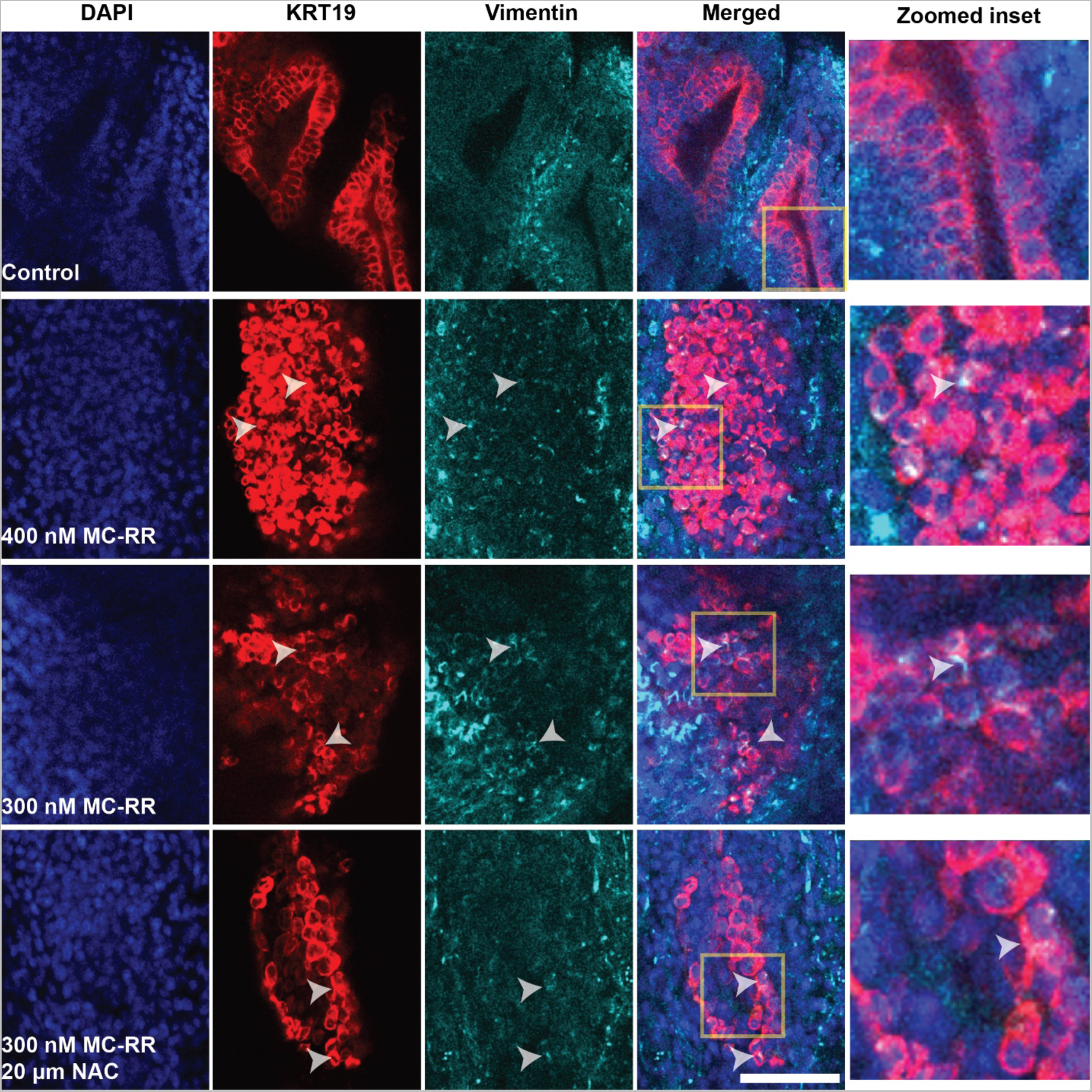
Vimentin expression in EHBDs treated with MC-RR and NAC. Representative images showing expression of vimentin in damaged cholangiocytes in EHBD explants in the presence of vehicle (control), 300 nM MC-RR, 400 nM MC-RR and 300 nM MC-RR+20 µM NAC for 24 h. White arrowheads highlight cells co-stained with KRT19 and vimentin. The last panel of each row shows the magnification of the yellow inset in the merged image. Cyan: vimentin, Red: KRT19, Blue: DAPI. All scale bars 50 µm.

**Supporting Figure S9:**
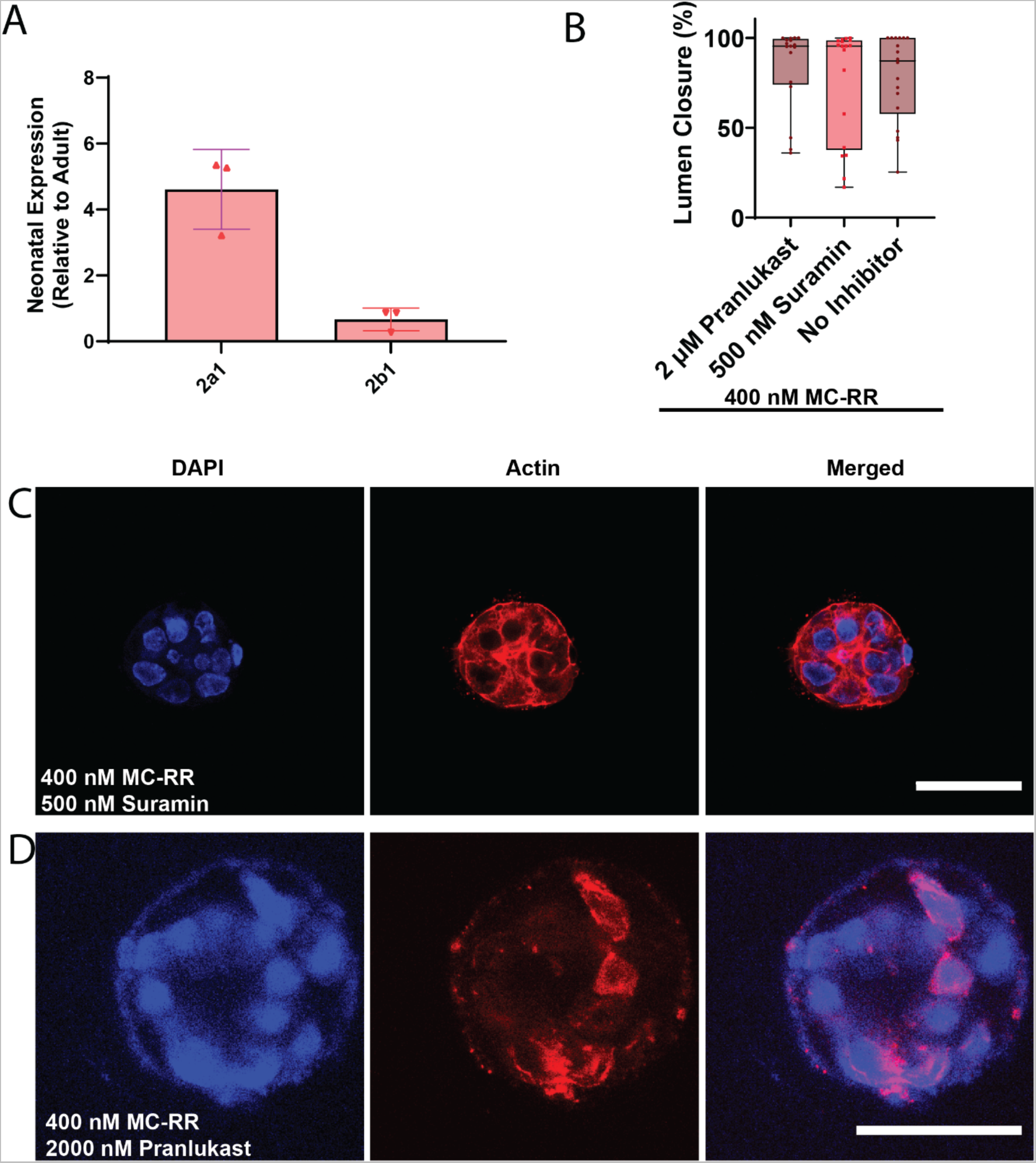
OATP inhibition does not rescue MC-RR-induced damage. A) Relative mRNA expression of OATP 2a1 and 2b1 in neonatal EHBD compared to adult EHBD. OATP 1a1, 1a4, 1a6 and 1b2 were below the detection limit in neonatal samples (n=3). B) Quantification showing no difference in MC-RR-induced lumen damage following OATP 2a1 inhibition using suramin and pranlukast (n=18 spheroids per condition from 3 independent experiments). C) Representative images showing neonatal spheroid damage in the presence of 400 nM MC-RR and 500 nM suramin for 24 h. D) Representative images showing neonatal spheroid damage in presence of 400 nM MC-RR and 2 µm pranlukast for 24 h. Red: actin, Blue: DAPI. All scale bars 50 µm.

**Supporting Figure S10:**
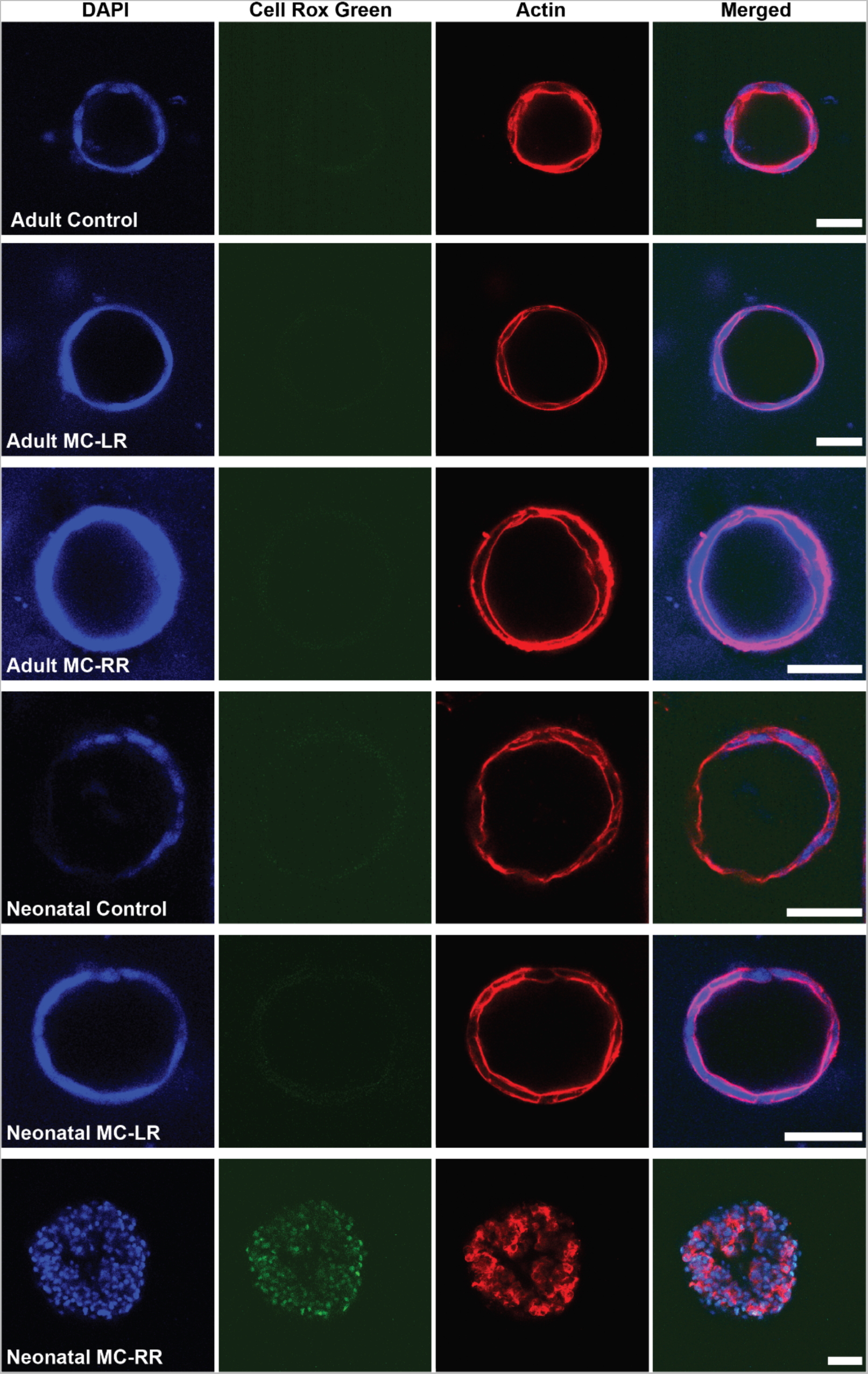
MC-RR specifically induces ROS in neonatal EHBD cholangiocyte spheroids. Representative images showing spheroids from adult and neonatal EHBD cholangiocytes treated with vehicle, 400 nM MC-LR and 400 nM MC-RR for 24 h. ROS were observed only in neonatal EHBD cholangiocytes treated with 400 nM MC-RR. Quantification in main Figure 6E. Red: actin, Green: Cell Rox Green, Blue: DAPI. All scale bars 50 µm.

**Supporting Figure S11:**
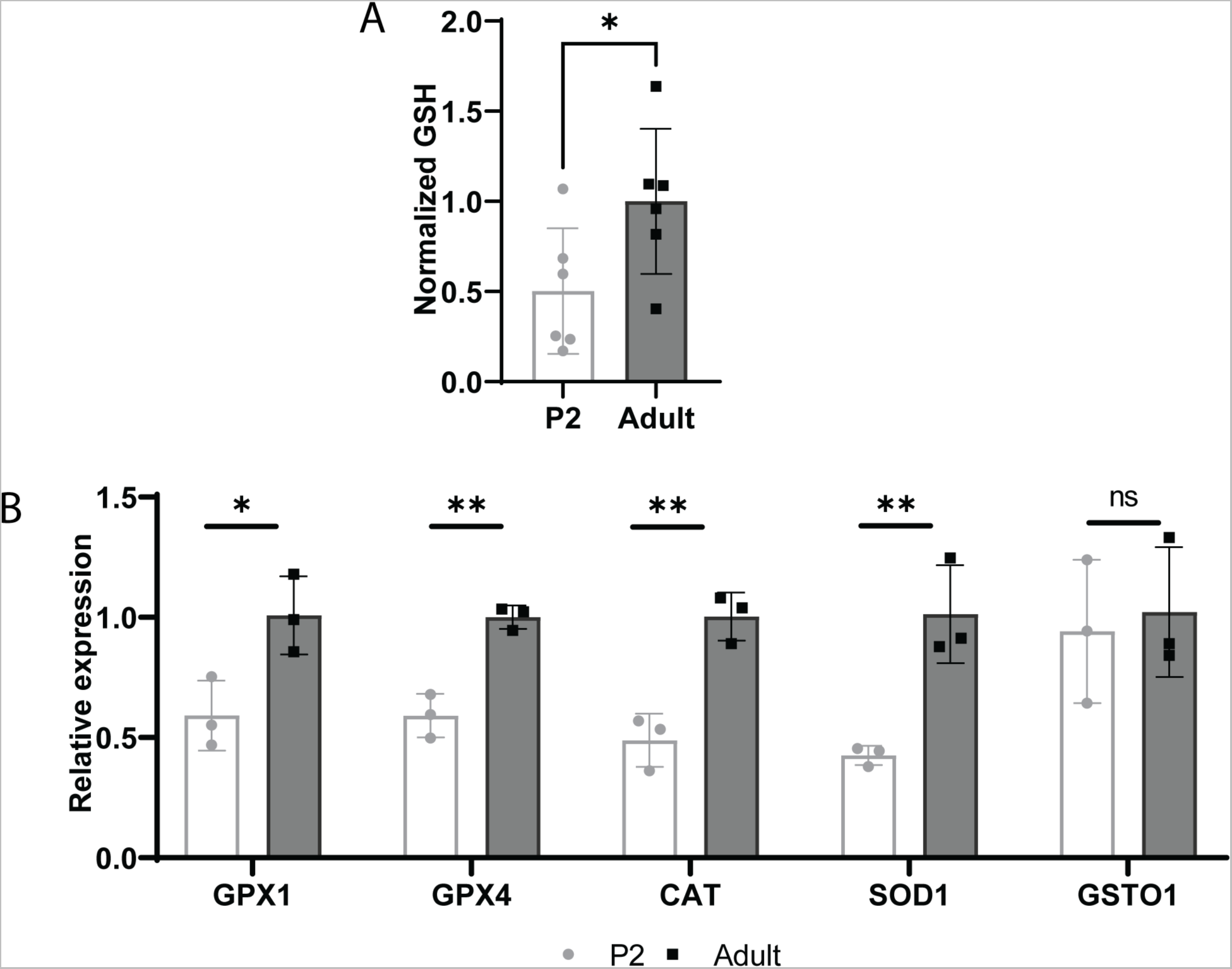
Neonates have immature antioxidant defense systems. A) GSH levels were quantified in protein samples obtained from P2 and adult mice, showing significantly lower GSH levels in neonates compared to adults (n=6). B) RNA samples from P2 and adult mice were analyzed for enzymes involved in antioxidant defense systems, demonstrating that all measured enzymes except GSTO-1 were significantly lower in neonates compared to adults (n=3).

**Supporting Figure S12:**
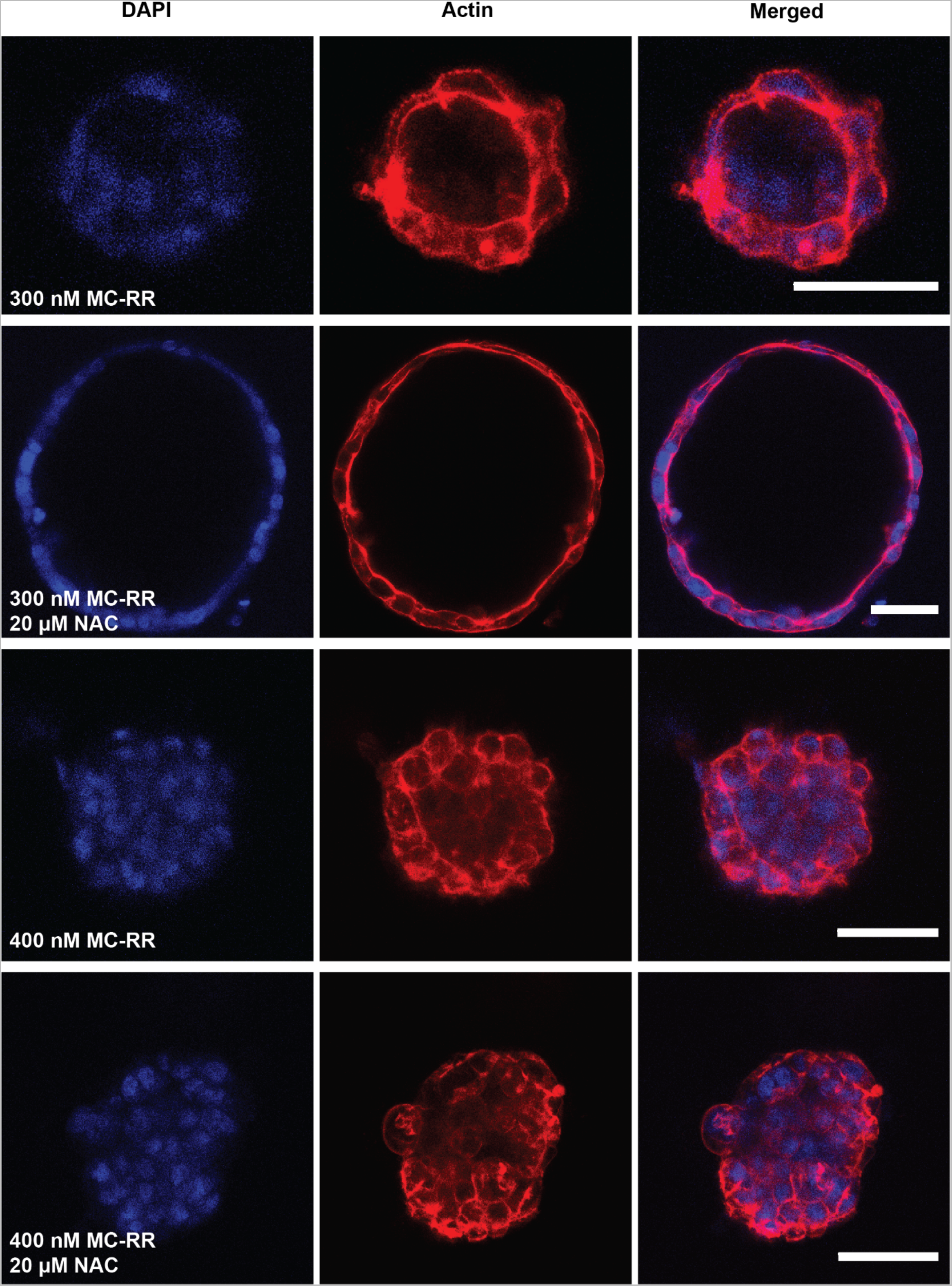
NAC rescues cholangiocyte damage in cholangiocytes treated with 300 nM MC-RR. Representative images showing spheroids treated with 300 and 400 nM MC-RR with and without 20 µM NAC for 24 h. Red: actin, Blue: DAPI. All scale bars 50 µm.

